# Cell-cycle dependent DNA repair and replication unifies patterns of chromosome instability

**DOI:** 10.1101/2024.01.03.574048

**Authors:** Bingxin Lu, Samuel Winnall, William Cross, Chris P. Barnes

## Abstract

Chromosomal instability (CIN) is pervasive in human tumours and often leads to structural or numerical chromosomal aberrations. Somatic structural variants (SVs) are intimately related to copy number alterations but the two types of variant are often studied independently. In addition, despite numerous studies on detecting various SV patterns, there are still no general quantitative models of SV generation. To address this issue, we develop a computational cell-cycle model for the generation of SVs from end-joining repair and replication after double strand break formation. Our model provides quantitative information on the relationship between breakage fusion bridge cycle, chromothripsis, seismic amplification, and extra-chromosomal circular DNA. Given single-cell whole-genome sequencing data, the model also allows us to infer important parameters in SV generation with Bayesian inference. Our quantitative framework unifies disparate genomic patterns resulted from CIN, provides a null mutational model for SV, and reveals new insights into the impact of genome rearrangement on tumour evolution.

## 1 Introduction

Chromosomal instability (CIN) is a major form of genome instability widely present in human tumours, which often leads to structural or numerical chromosomal aberrations and plays an important role in cancer evolution [1, 2]. Somatic structural variants (SVs) are large genomic rearrangements showing a variety of patterns and have been classified into different types, such as duplication, deletion, inversion, translocation, and other complex variants [3–10]. Duplication and deletion are also called copy number alterations (CNAs), which include whole genome doubling (WGD) as well and are often studied separately from SVs [7, 9, 11, 12]. Typical complex SVs include breakage-fusion-bridge (BFB) cycle [13], chromothripsis [14], extrachromosomal circular DNA (ecDNA) [15], seismic amplification [8], and chromoplexy [16]. From these various patterns, it is important to investigate the underlying mechanisms, which will provide further insights into CIN and cancer evolution and potentially inform treatment [5, 17–21].

Each SV manifests as the spatial apposition of breakpoints [17] that is caused by the interplay of DNA damage, replication errors, and repair pathways occurring throughout the cell cycle [22]. Many DNA damage events involve double strand breaks (DSBs), which can be repaired by two broad mechanisms: homologous recombination (HR), which has high fidelity but requires extensive sequence homology at the breakpoints; or non-homologous end joining (NHEJ) that needs no sequence homology but is error-prone [6, 23, 24]. Indeed, NHEJ is known to play a major role in the formation of focal deletions and translocations [25], chromothripsis [26–30], ecDNAs [15], and various simple and complex SVs after chromosome segregation errors [31]. How erroneous DNA repair mechanisms directly contribute to the complex SVs observed in cancer genomes still remains unclear [21, 24].

This question cannot be answered by experimental approaches alone due to technical limitations [20, 32, 33] and requires quantitative models which can encode our knowledge of the underlying biological processes. The quantitative models enable long-term simulation to study evolutionary processes inaccessible by experiments and provide a basis for statistical inference. A few models have been developed to investigate CNAs [2, 33–36] and certain types of SVs, such as chromothripsis [14, 37], chromoplexy [16], ecDNA [15, 16], neochromosome [38], and seismic amplification [8]. However, these models focused on specific SV types and local genomic regions, therefore missing the underlying processes that relate all of these variant classes. This is like studying wind speed, precipitation, thunderstorms, and tornadoes without understanding changes in air temperature and air pressure.

To address this gap, we developed a conceptually simple, yet unifying framework to model the dynamics of SV formation. Our multi-scale computational model consists of simulated cells progressing through cell cycles via a stochastic birth-death branching process where fitness effects can be captured. A cell undergoes random DSB repair through end-joining and genome replication in interphase and then correctly or incorrectly divides in mitosis. Using our model, we provide direct quantitative evidence that different types of SVs are generated during the same ongoing DNA repair and replication process, including BFB cycle, chromothripsis, ecDNA, and seismic amplification. We then apply the model with Bayesian inference techniques to infer important parameters in single-cell whole-genome sequencing data, such as DSB rate per cycle, probability of WGD per cell, and otherwise inaccessible information including mean number of chromosomal fusions per cycle and mean number of ecDNAs per cell.

## 2 Results

### A stochastic model of SV generation across cell cycles

To model SV generation within a cell, we introduced DSBs and repaired them with random end-joining across the cell cycle. Cells divided according to a stochastic birth-death branching process (Fig. 1a, Table S1, Methods). For simplicity, we did not explicitly consider sequence homology at breakpoints when repairing DSBs, and hence the erroneous repair in our model is considered as mainly deriving from NHEJ, which was previously shown to generate many SVs in cancer genomes [6]. Some SVs such as tandem duplications may be generated by homologous repair deficiency (HRD) or DNA replication errors [25, 39]. However, since only around 12% of cancers show tandem duplication phenotype [40], we did not include these processes in our model.

**Fig. 1.**
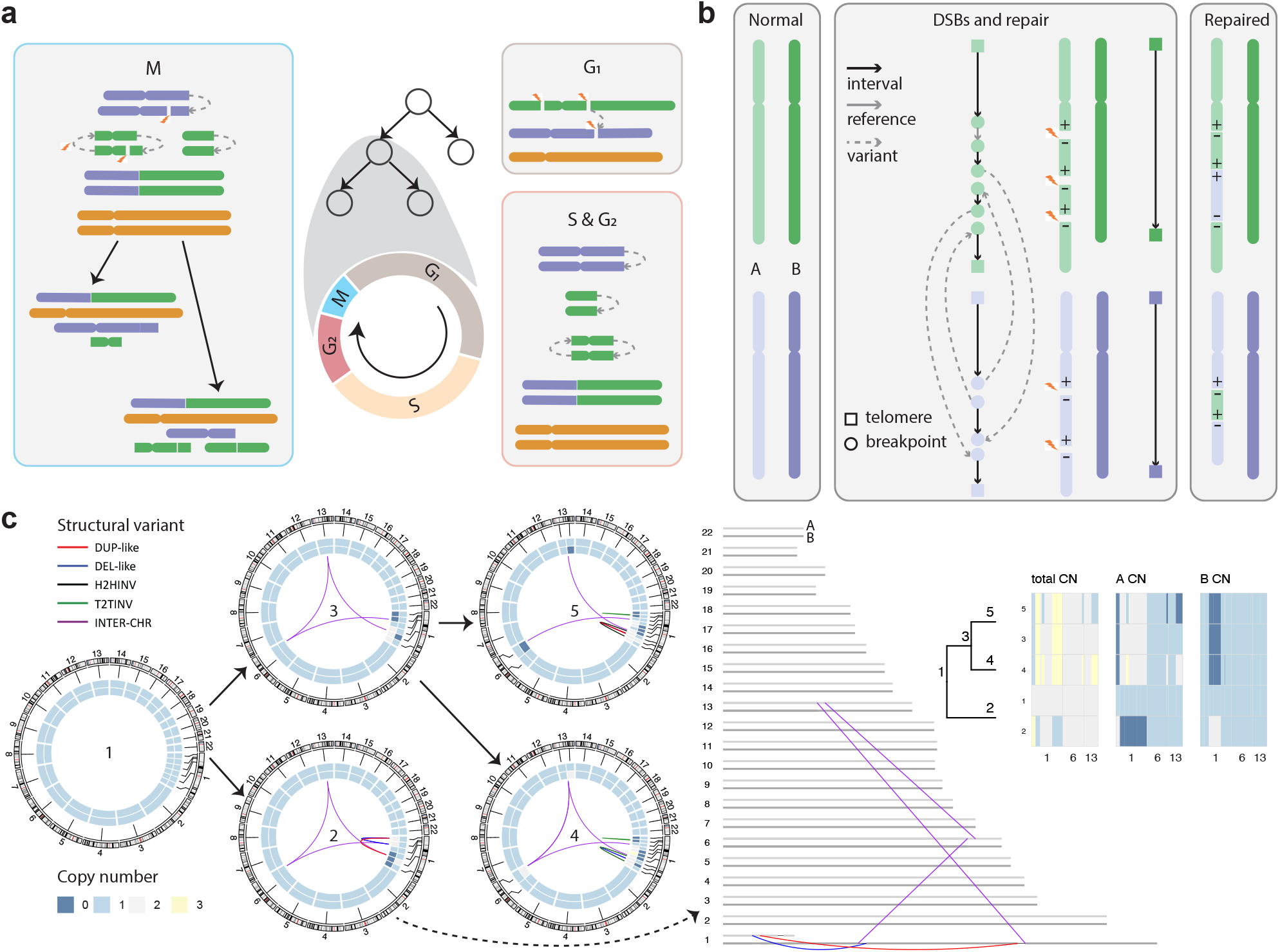
The stochastic cell-cycle model of SV generation from DNA repair and replication. **a**, Illustration of the cell-cycle model along a branching process. The cell cycle contains interphase and mitosis (*M* phase), where interphase consists of DNA replication with three phases: *G*_1_ (growth) in which the cell grows, *S* (synthesis) in which the cell replicates DNA, and *G*_2_ (growth) in which the cell grows to prepare for mitosis. In *G*_1_, the joining of the broken green and purple chromosomes forms chromosome fusion. In *S* and *G*_2_, after DNA replication, the joining of *p* arms of the green chromosome and *q* arms of the purple chromosome forms sister chromatid fusion respectively, whereas the joining of the remaining parts of the green chromosome forms a circular DNA. In *M*, the two fused chromosomes with more than one centromeres are randomly broken and distributed to two daughter cells, imitating BFB. At the same time, the two copies of the chromosomes with one centromere are distributed to each daughter cell, and the green chromosome without centromere is randomly distributed to a daughter cell. **b**, The graph representation of a diploid genome. Left, two pairs of normal chromosomes with light colour indicating homolog A and dark colour indicating homolog B. Middle, one homolog of each chromosome is broken at some breakpoints and rejoined randomly. The normal chromosome is represented by two telomeres connected with an interval edge. The broken and repaired chromosome is represented by the joining of both telomeres and breakpoints with interval, reference, and variant edges. Right, the repaired chromosomes with SVs (translocation and inversion). **c**, The accumulation of SVs in two cell cycles starting from a normal cell. Left, the circos plots of all the cells with cell lineage information, where the inner and outer heatmaps within the circle represent copy numbers of homolog A and B respectively. Middle, the haplotype graph of cell 2, where the breakpoints are located at different homologs which are not distinguished in the circos plots. Right, the cell lineage tree and copy numbers of chromosomes with CNAs. The copy numbers in the two daughter cells after a cell cycle are always reciprocal. DUP-like: tandem duplication-like patterns, DEL-like: deletion-like patterns, H2HINV: head-to-head inversion, T2TINV: tail-to-tail inversion, INTER-CHR: inter-chromosomal SV.

We also assumed that a fraction of DSBs may remain unrepaired due to checkpoint deregulation such as TP53 mutation [22]. As SVs may bring selection advantages or disadvantages to cancer cells, we incorporated a simple model of selection based on the density of oncogenes (OGs) and tumour suppressor genes (TSGs) [2, 41], in which karyotypes with a higher oncogenic propensity were favoured.

To keep track of breakpoint joining on the complete genome that becomes more shattered with additional DSBs across cell cycles, we used a diploid interval adjacency graph *G* = (*V, E*) to represent each genome [42], where *V* represents the set of breakpoints (nodes) and *E* represents their adjacencies (edges) (Fig. 1b). There are three types of adjacencies according to the positions in the reference genome, including an interval adjacency between two endpoints of a genomic interval, a reference adjacency connecting two adjacent intervals, and a variant adjacency caused by SVs which connects two non-adjacent intervals. By using a graph representation to capture the complexity of SVs in a complete genome, we can generate realistic data including CNAs and SVs for all the cells in the final population.

To illustrate our model in simulating SV formation over time, we show the accumulation of SVs in two cell cycles starting from a normal cell with intuitive plots (Methods, Fig. 1c). The formation of SVs is clearly shown in a step-wise way at the haplotype level, helping to understand the link between DNA repair and replication with different types of SVs.

### The simple repair process model explains the formation of complex SVs

With appropriate parameters, our model enables forward simulations which generate realistic inter-chromosomal and intra-chromosomal rearrangements, such as BFB cycle, chromothripsis, focal and seismic amplification, ecDNA, chromoplexy, and parallel haplotype-specific CNA [20, 21, 43] (Methods).

In our model, the fusion of sister chromatids of a telomere-lacking chromosome in interphase leads to the forming of a chromosome bridge and the asymmetric bridge breakage in mitosis (Fig. 1a), which was shown to generate either simple breaks or local fragmentation that trigger iterative cycles of complex mutational events including chromothripsis interwoven with BFB cycles [20]. To validate this phenomenon, we introduced one unrepaired DSB on a random chromosome followed by either simple breaks or local fragmentation in the simulations. The simulation with simple breaks recaptured the typical features of a BFB cycle, including staircase-like focal inverted duplications or fold back inversions (FBIs) with clustered breakpoints adjacent to terminal losses or gains on the same homolog [21, 43] (Fig. 2a, b). Under selection, cells with higher-level amplifications seem slightly favoured, which is likely due to a few cells with higher copy numbers of OGs such as CARD11 and EGFR (Table S2). The results also directly demonstrate that an ongoing BFB cycle initiated by the formation of a chromosome bridge caused progressive shorting of a homolog and serrate structural variation (SSV), which were recently reported in bulk or single-cell whole-genome sequencing data and suggested to reflect ongoing BFB cycles or mutational process [20, 43] (Fig. 2a). With local fragmentation, the random segregation of highly broken dicentric chromosomes led to a number of chromothripsis, ecDNAs, and seismic amplifications which started appearing at similar times [21] (Fig. 2d, e, Table S3). The sizes of simulated ecDNAs mainly ranged from hundreds of bases to several megabases, consistent with those reported in literature [44], and there were slightly more different types of ecDNAs per cell under positive selection (Fig. A1). Due to weak selection forces (Fig. A1, Table S4), the numbers of complex SVs generated under neutral evolution and selection were similar. Our model is the first to show that BFB cycles, chromothripsis, and ecDNAs are closely related by erroneous DNA repair (likely from NHEJ) and replication at the whole genome level in silico, consistent with the known experimental discovery that BFB cycles and chromothripsis trigger ecDNA formation and amplification along with NHEJ inhibitors [15]. Our model also suggests that chromothripsis can contribute to seismic amplification, which agrees with previous work that shows seismic amplification can be formed via chromothripsis followed by circular BFB or recombination of ecDNAs [8].

**Fig. 2.**
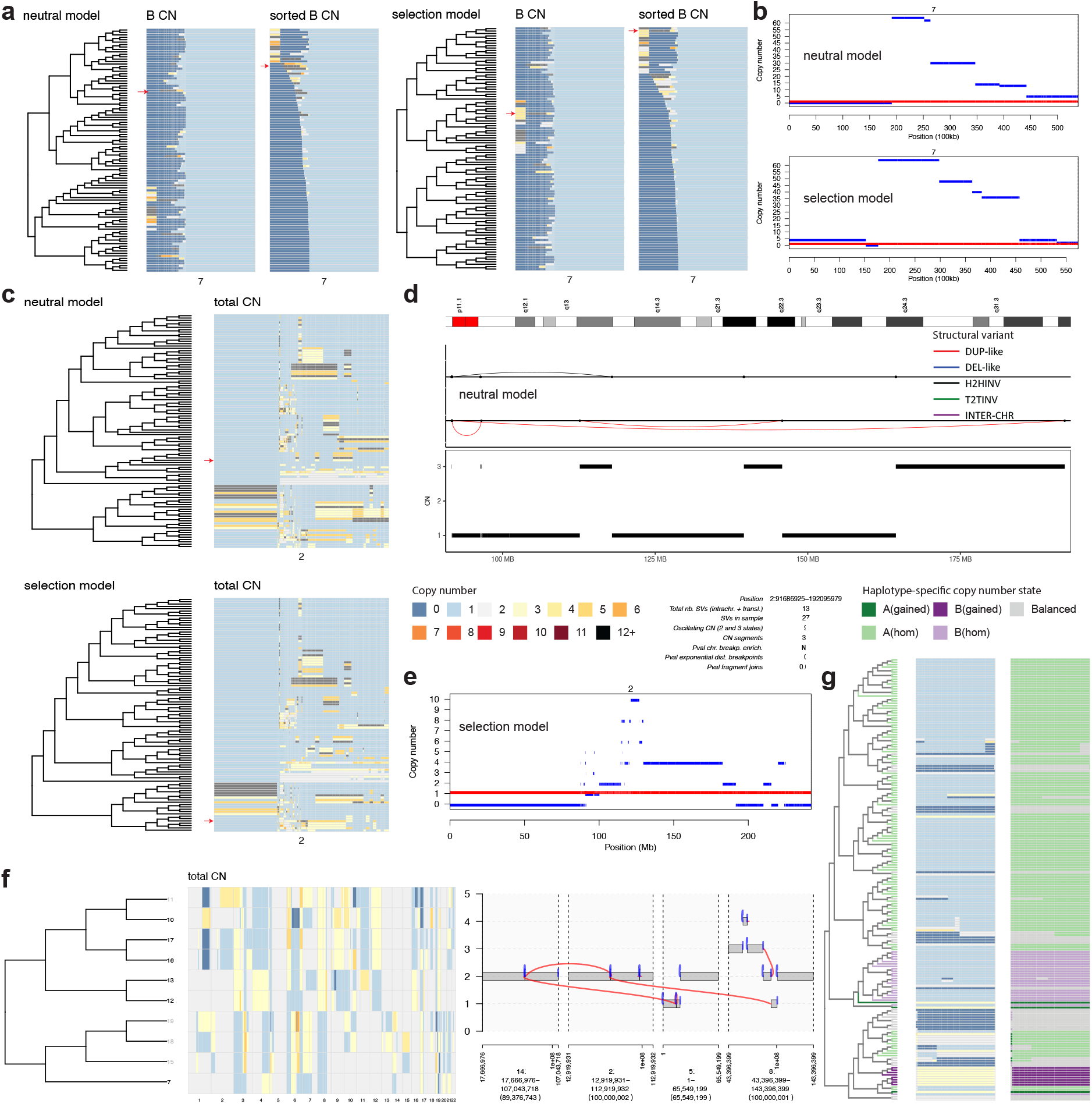
Demonstration of SV patterns generated by the cell-cycle model. **a**, The cell lineage tree of 100 cells and their corresponding copy numbers on chr7 from the simulation with simple breaks under neutral evolution and selection, where the heat map of homolog B on the right is sorted to show SSVs. **b**, The haplotype-specific CNAs of two cells (indicated by red arrows in **a**) suggest staircase-like focal amplifications. **c**, The cell lineage tree of 100 cells and their corresponding copy numbers on chr2 from the simulation with local fragmentation under neutral evolution and selection. **d**, Chromothripsis detected in one cell after the eighth cell cycle under neutral evolution (indicated by top red arrow in **c**). **e**, Haplotype-specific CNAs of one cell at the fifth cell cycle with seismic amplification under selection (indicated by the bottom red arrow in **c**). **f**, The cell lineage tree of 10 cells and their corresponding copy numbers across all chromosomes from the simulation with multiple misrepaired DSBs, as well as a detected chromoplexy in cell 10. The cells without chromoplexy are shown in grey. **g**, The cell lineage tree of 183 cells (excluding 17 cells lacking the pattern) and their corresponding total copy numbers and haplotype-specific copy number states at region (19444803, 27298459) of chr22.

Balanced rearrangements such as chromoplexy were hypothesised to arise from incorrect rejoining of several simultaneous DSBs in multiple chromosomes, which may span several successive events [6, 16, 19]. To validate this hypothesis, we introduced a few misrepaired DSBs on several chromosomes in the simulation, which showed that chromoplexy appeared after just a few cell cycles (Fig. 2f, Fig. A2). The underlying processes of chromoplexy are still largely unknown [21], and experimental studies have recently been done to investigate whether co-localization of multiple DSBs can stimulate chained inter-chromosomal and intra-chromosomal translocations typical for chromoplexy [45]. This is now confirmed in our simulations which may help to further understand the processes generating chromoplexy by fitting experimental data in the future. As a DSB occurred on one homolog of the affected chromosome, we also observed parallel evolution of haplotype-specific CNAs that affect different alleles of the same sites in different cells when we continued the simulation until a larger number of cells (Fig. 2g). Parallel haplotype-specific CNAs have been commonly detected in cancer genomes, which may influence transcription and make it challenging to compute variant allele frequency in bulk sequencing data [2, 43]. Our model provides a straightforward way to simulate them and may further contribute to understanding their roles in cancer evolution.

### Role of cell cycle in formation of complex SVs including chromothripsis

We explored how DSB rate per cycle, percentage of unrepaired DSBs per cycle, and scale of local fragmentation (measured by mean number of DSBs per fragmentation) affected the formation of chromothripsis, which was linked with poor survival or prognosis in cancer patients [46] (Methods). The results suggest that chromothripsis events were generated in one or two cell cycles when multiple DSBs co-located on chromosomes with all of them being misrepaired or a fraction remaining unrepaired (Fig. 3). Even with a larger number of DSBs in one cell cycle, fewer chromothripsis events occurred, along with no seismic amplification as well as a smaller number of chromosome fusions (BFB cycles) and ecDNAs, which all increased in two cell cycles. Therefore, chromothripsis seems more likely to be formed in a cascade of mutational events as demonstrated in recent experiments [20] rather than in a one-off event. The numbers of BFB cycles and ecDNAs generally increased with the number of cell cycles, the DSB rate, the scale of local fragmentation, and the percentage of unrepaired DSBs, whereas the numbers of chromothripsis and seismic amplifications fluctuated sometimes as a result of the strict criteria used in detecting these events. When higher levels of local fragmentation were introduced, the involved chromosome was shattered into more pieces and hence generally led to more BFB cycles, chromothripsis, ecDNAs, and seismic amplifications, which confirms the experimental results that most SVs occurred after local fragmentation [20]. With high-level local fragmentation, the numbers of BFB cycles, chromothripsis, and ecDNAs under two cell cycles were much larger than those under one cell cycle, especially when DSB rate and percentage of unrepaired DSBs were also higher (Fig. A3). However, the differences under the two cycles with or without consecutive interphase DSBs were relatively small, suggesting that the high probability of generating chromothripsis in two cell cycles with just a higher number of unrepaired interphase DSBs and local fragmentation in one cell cycle, which may help to explain the wide prevalence of chromothripsis and ecDNAs in cancer genomes [27, 28, 47].

**Fig. 3.**
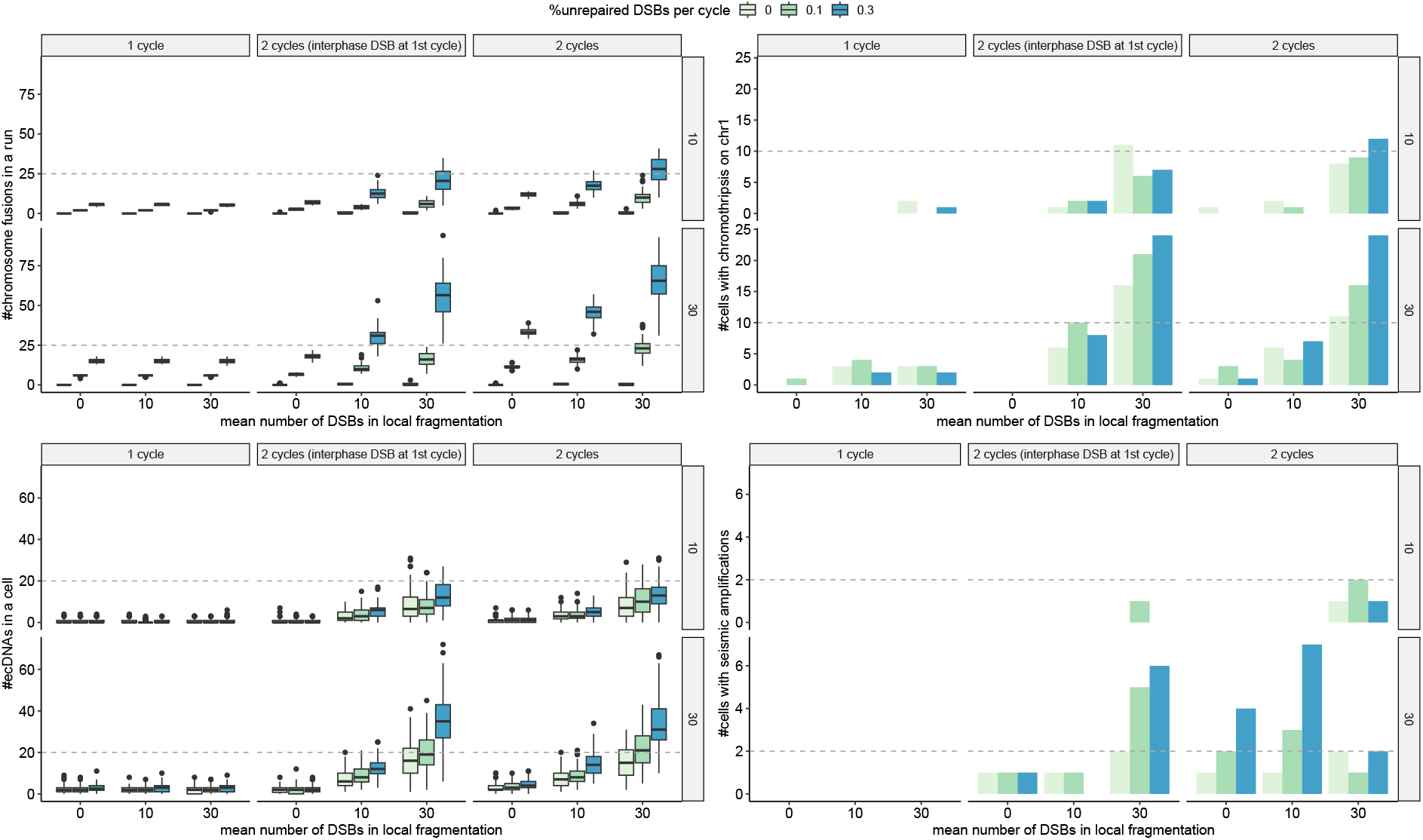
The distributions of complex SVs under different parameter settings in one or two cell cycles. The labels at the right of each plot indicate DSB rates per cycle. For each run, we consider the two cells at the end of either the first or second cycle depending on the number of cycles. The box plots show the median (centre), 1st (lower hinge), and 3rd (upper hinge) quartiles of the data; the whiskers extend to 1.5 times of the interquartile range (distance between the 1st and 3rd quartiles); data beyond the interquartile range are plotted individually.

### Exploration of long-term evolutionary dynamics of ecDNA

Using simulations of larger cell populations under both neutral evolution and selection, our model can help to reveal the evolutionary dynamics of SVs in the long run. The ecDNA-based oncogene amplifications are common and associated with poor outcome across multiple cancer types [48], whose evolutionary dynamics has been recently well studied [49, 50]. To compare with previous results, we continued our simulation of one unrepaired DSB with local fragmentation until 1000, 3000, and 5000 cells to see the long-term effect of selection on ecDNAs. As we simulated ecDNAs from DSB repair in the background of other SVs rather than in isolation, there may be different types of ecDNA with various numbers of copies in a cell. This is more realistic when compared to previous studies which focused on a single type of ecDNA with a specific oncogene [49, 51], since multiple types of ecDNA have been detected in 26 out of 83 patient samples with ecDNAs [50].

Our results confirm that ecDNAs show extreme variation across cells and increasing copy numbers and complexity over time [49] (Fig. 4). The numbers of chromosome fusions under selection were much smaller than those under neutral evolution, which suggests that many fusions were disadvantageous, as more cells were under negative selection than positive selection (Fig. 4a). Although the numbers of positively selected cells were relatively small, the fraction of cells with ecDNAs under positive selection kept increasing with rising fitness, whereas those under neutral evolution and negative selection kept decreasing, which is consistent with previous mathematical analysis (Fig. 4a, Fig. A4) [49]. For a specific type of ecDNA, the distribution of ecDNA copy numbers under strong positive selection was shown to shift towards higher copy numbers over time [49], which is confirmed in the copy number distribution of the most common ecDNA type in our simulations (Fig. 4b). Although the sizes of ecDNAs and the mean ecDNA copy numbers per cell remained similar under both neutral evolution and selection, the numbers of ecDNA types and the maximum ecDNA copy numbers per cell were slightly higher under positive selection and neutral evolution (Fig. 4c). The slight changes of ecDNA sizes, moderate increases of mean copy numbers, and larger increases of maximum copy numbers over time (Fig. 4c) are consistent with previous discoveries obtained from patients with Barrett’s oesophagus, which showed ecDNA sizes were not significantly different whereas ecDNA copy numbers became significantly higher during malignant progression [50].

**Fig. 4.**
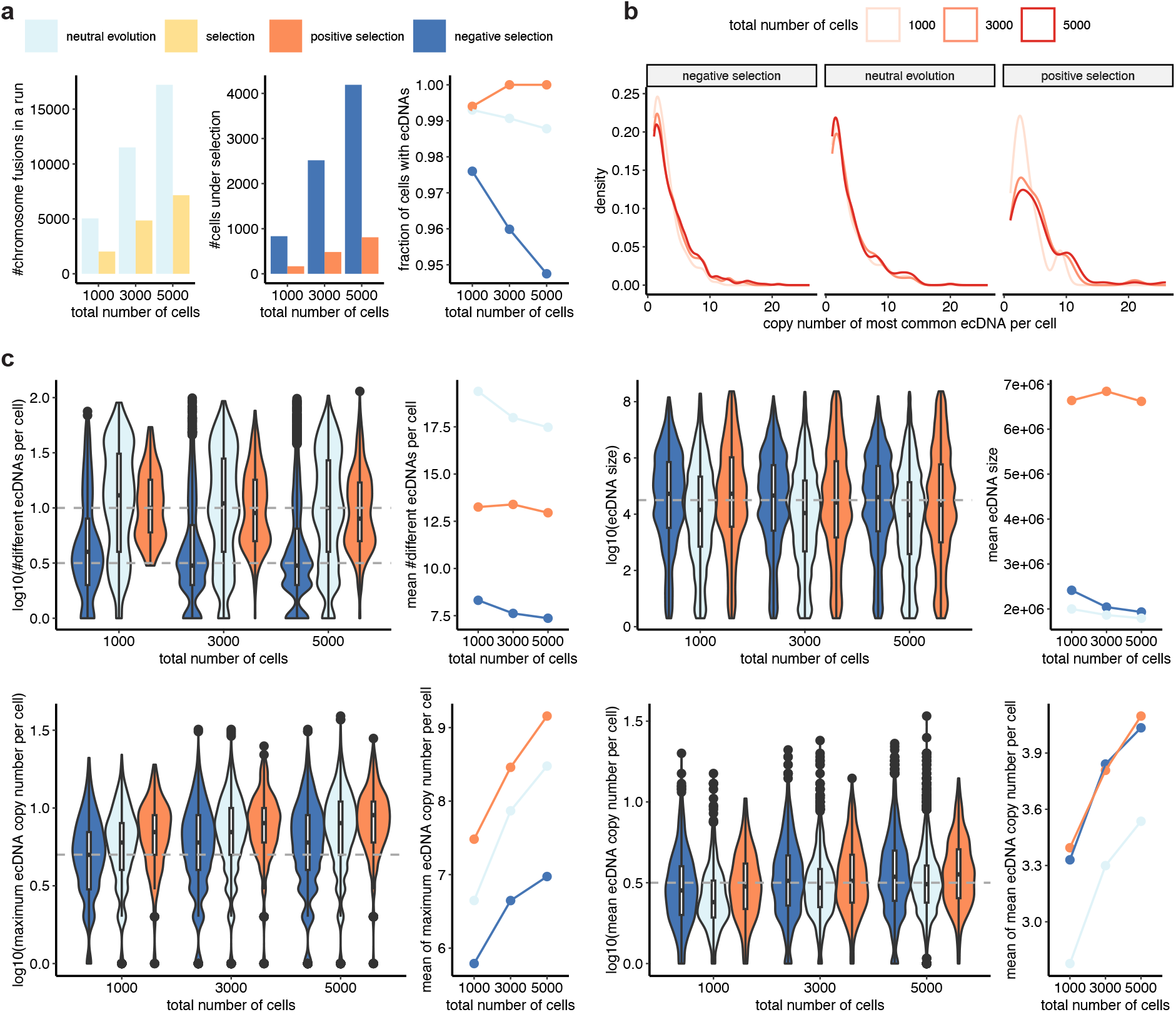
The evolutionary dynamics of ecDNAs under different selection forces over time. **a**, The numbers of chromosome fusions and cells with ecDNAs. **b**, The distributions of copy numbers of the most common ecDNA type. **c**, The distributions of the number, size, maximum copy number, and mean copy number of different ecDNAs per cell under different selection forces. The box plots show the median (centre), 1st (lower hinge), and 3rd (upper hinge) quartiles of the data; the whiskers extend to 1.5 times of the interquartile range (distance between the 1st and 3rd quartiles); data beyond the interquartile range are plotted individually.

### Validation of parameter inference on simulated data

To gain insight into the formation and evolution of rearranged genomes, we sought to develop a simulation-based inference approach using approximate Bayesian computation (ABC), which can be applied when the likelihood function is intractable [52]. We first came up with a set of informative and low-dimensional summary statistics based on domain knowledge, which aggregated over the complexity of genome rearrangement events (Table S5). We inferred the posterior distributions of three key parameters in SV formation with ABC sequential Monte Carlo (SMC): DSB rate per cycle, fraction of unrepaired DSBs per cycle, and probability of WGD per cell, whereas the values of the other parameters of less interest were fixed to default values (Table S1, Methods). We checked the predictive power of our model by generating the posterior predictive distributions of two summary statistics arising in the simulation but often not directly observable from genome sequencing data: mean number of chromosome fusions per cycle and mean number of ecDNAs per cell.

The results suggest that both DSB rate and fraction of unrepaired DSBs were accurately estimated especially when real values were higher, with posterior medians close to real values (Fig. S1 and S2), although the distributions of posterior means suggest that they were slightly overestimated (Fig. 5a,b). The posterior means of the probability of WGD were more overestimated (Fig. 5c), and it is harder to accurately infer large values, as reflected by the wider posterior distributions (Fig. S1 and S2). This is because a large probability of WGD is very likely to give rise to repeated WGD whereas our simulation is constrained to at most one WGD per cell. The mean number of fusions per cycle was generally underestimated yet improved in precision with the increase of the real fraction of unrepaired DSBs, as chromosome fusions are mainly caused by broken ends. The mean number of ecDNAs per cell was slightly overestimated and became more accurate when there were more ecDNAs in the data (Fig. 5d,e, Fig. S1 and S2). The obtained summary statistics from posterior predictive distributions were generally consistent with those observed in the simulated truth, whereas the deviances of some statistics were slightly larger, likely due to the difficulty in fitting a large number of summary statistics (Fig. S3). Taken together, these results show that our model-based simulation allows reliable Bayesian inference of important parameters in SV formation when there is sufficient information in the data and hence can be used to understand the processes underlying real genome rearrangments.

**Fig. 5.**
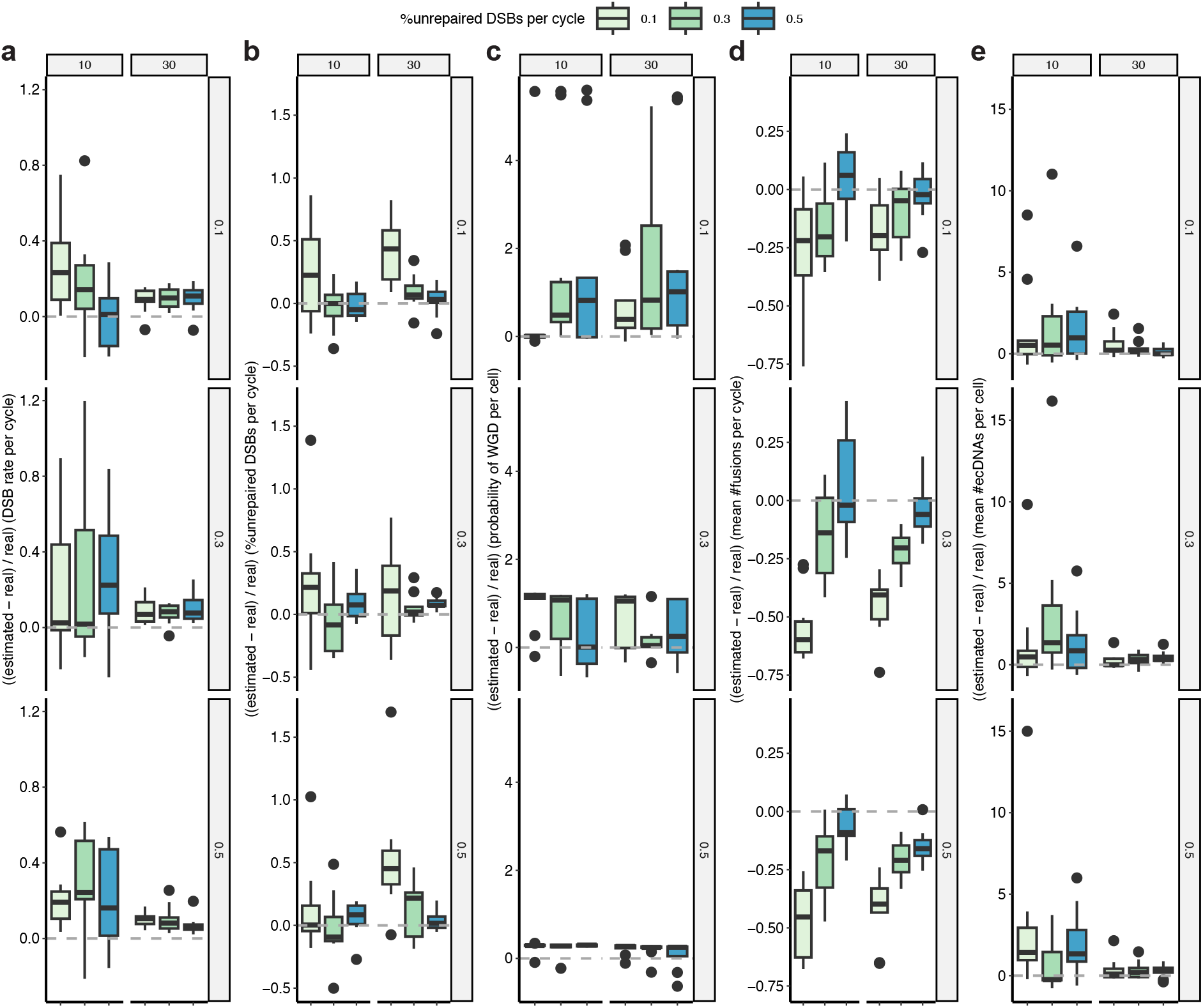
The accuracy of parameter inference with ABC SMC on simulated data. The proportion of differences between estimated and real values of the three inferred parameters (**a**,**b**,**c**) and two summary statistics (**d**,**e**). The box plots show the median (centre), 1st (lower hinge), and 3rd (upper hinge) quartiles of the data; the whiskers extend to 1.5 times of the interquartile range (distance between the 1st and 3rd quartiles); data beyond the interquartile range are plotted individually. There are 10 data points for each box plot, which represent the posterior means of 10 runs under the same parameter setting. The labels at the top of each plot show DSBs rate per cycle whereas the labels at the right show probabilities of WGD per cell.

### Fitting the model to single-cell whole-genome sequencing data of epithelial cell lines and patient-derived xenografts

To demonstrate the utility of our model in learning SV generation mechanisms, we analyzed 20 single-cell whole-genome sequencing datasets [43], where 8 datasets are from 184-hTERT mammary epithelial cell lines, and 12 datasets belong to FBI tumours from patient-derived xenografts (PDXs) of primary cancer patients, which are enriched in CCNE1 amplifications (Table S6). As the coverage of single-cell data was too low to detect SVs, pseudobulk data were used to detect breakpoints in individual clones of each dataset. Although this means we cannot apply our model to individual cells, we can still gain insights from clone-specific data where typical patterns of SVs can be identified. We simulated the branching process until it reached the number of clones in each dataset, and parameter interpretations were adjusted accordingly to clone expansion rather than cell division.

The results (Fig. 6a) were largely consistent with the observed SV patterns (Fig. S4) and showed high heterogeneity of key parameters across different datasets. Uniquely, our inference gave rise to posterior predictive distributions suggesting potential bounds on the unobserved numbers of BFB cycles (chromosome fusions) and ecDNAs. For a few datasets, the inferred DSB rates were more different from empirical DSB rates (Fig. A5a), which could be due to SVs generated from mechanisms not captured in our model such as HRD or replication errors [25, 39]. On datasets with fewer polyploid clones, the inferred probability of WGD per cell was similar to the fraction of polyploid clones in each sample. In contrast, on datasets with more polyploid clones, the inferred value was generally smaller than the real fraction due to the constraint of at most one WGD in the simulation (Fig. A5b). We detected few chromosome fusions and ecDNAs in the cell line datasets (Fig. 6a),which generally have fewer SVs than the FBI tumour PDX datasets. Our predictions indicate that SA1188, on average, has around one fusion and no ecDNA, which agrees with the typical patterns of BFBs detected on chr3q [43]. As it is still challenging to detect ecDNAs from single-cell data [53], our inferences may provide hints for further validations.

**Fig. 6.**
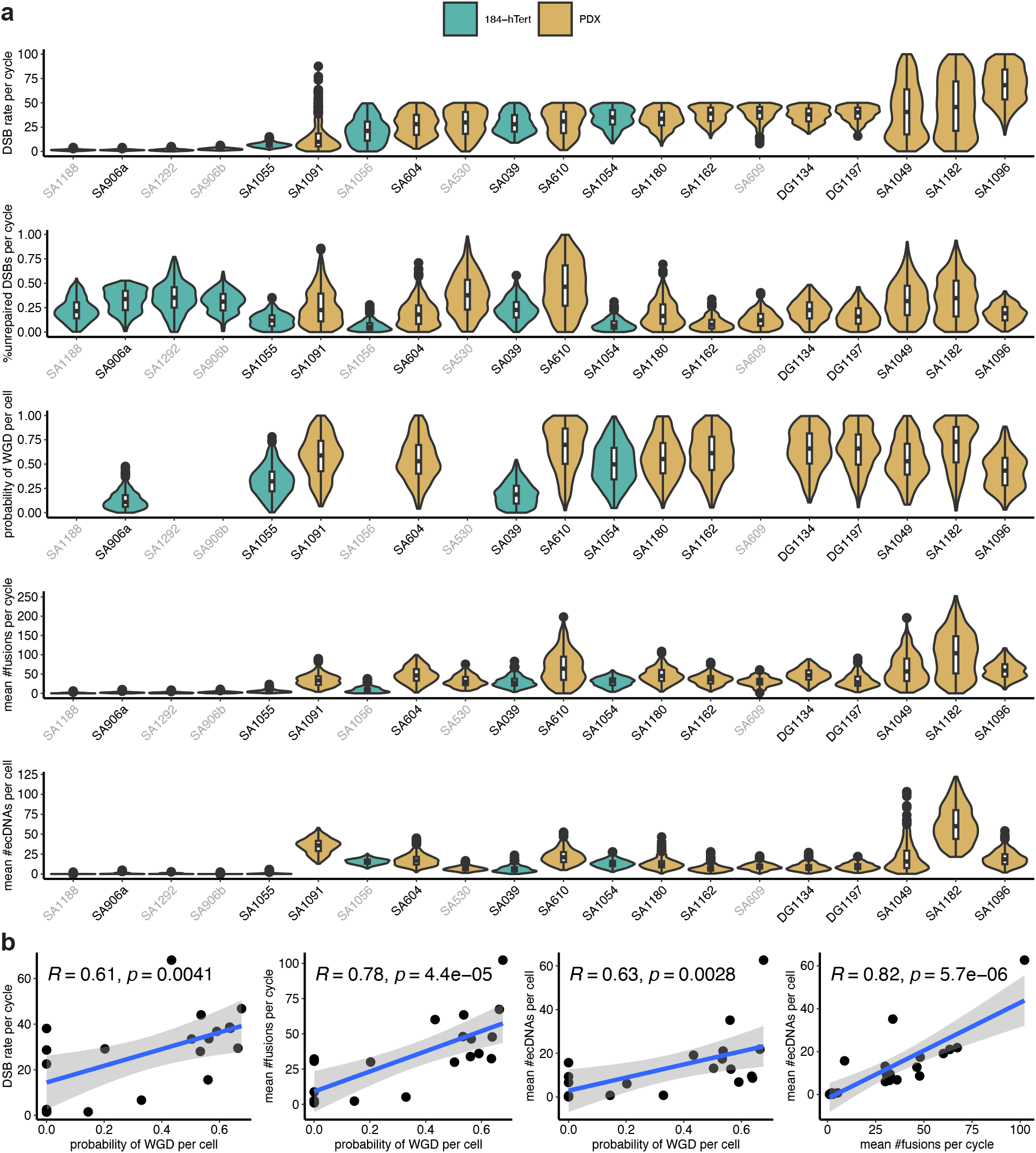
Results of fitting the cell-cycle model to single-cell whole-genome sequencing data with ABC SMC. **a**, The posterior distributions of inferred parameters and the posterior predictive distributions of the unobserved numbers of chromosome fusions and ecDNAs. The names of datasets with WGD are shown in grey. The box plots show the median (centre), 1st (lower hinge), and 3rd (upper hinge) quartiles of the data; the whiskers extend to 1.5 times of the interquartile range (distance between the 1st and 3rd quartiles); data beyond the interquartile range are plotted individually. **b**, The pairwise correlations among the inferred DSB rate per cycle, probability of WGD per cell, mean number of fusions per cycle, and mean number of ecDNAs per cell. The Spearman correlation coefficient and corresponding p-value are shown for each plot. The shaded area shows the 95% confidence interval of linear regression. Each point represents the posterior mean.

To understand the relationship among inferred parameters, we computed their posterior means for each dataset and plotted their pairwise correlations (Fig. 6b, Fig. A6). We found significant correlations between chromosome fusions and ecDNAs in both single-cell whole-genome sequencing data and our simulated data (Fig. A7), suggesting that ecDNAs were mainly derived from BFB cycles and our model captured this feature. The posterior means of DSB rate, mean number of chromosome fusions, and mean number of ecDNAs significantly increased with probability of WGD (Fig. 6b). This is an emergent property of the dynamics since the model doesn’t contain these dependencies, as demonstrated in the simulation results (Fig. A7a-c). Thus, we have shown that WGD can affect the generation of SVs through simple cellular processes. The significant correlation between the probability of WGD and DSB rate is consistent with previous reports on increased levels of CNAs in cancers with WGD [11], elevated CNA rates after WGD in breast tumours [35], and rapid accumulation of arm-level CNAs triggered by WGD in Barrett’s esophagus and esophageal adenocarcinoma [54]. There have been no previous results on the correlations between WGD and chromosome fusions or ecDNAs, which are only implicated in previous studies showing that WGD correlated with CCNE1 amplifications [11], which is frequently targeted by BFB cycles [7] that can lead to chromothripsis and ecDNAs [15]. Moreover, the previous observation that chromothripsis occurred after WGD in 74% of 194 cases where event orders can be distinguished [27] also implies the correlation between WGD and ecDNAs, as chromothripsis is a major driver of ecDNA formation and amplification [15]. Compared with cell line data, FBI tumours seem more likely to undergo WGD and have a much higher DSB rate, a slightly larger fraction of unrepaired DSBs, and much more BFB cycles and ecDNAs (Fig. A8). The correlations of WGD with DSB rate, chromosome fusions, and ecDNAs may be explained by the enrichment of CCNE1 amplification in FBI tumours. Given that CCNE1 amplification is also associated with large tandem duplications [40], it seems instrumental in SV formation and CIN.

## 3 Discussion

In summary, we proposed a quantitative cell-cycle model for the generation of a wide spectrum of patterns caused by CIN at the whole genome level. Our model incorporates DNA damage along with erroneous repair and replication processes, which has not been attempted previously. This approach provides a natural way to integrate CNAs and SVs, which have often been analysed independently despite being intrinsically coupled by unifying mutational processes [7]. Through simulations under diverse scenarios, we showed how various chromosomal alterations developed and correlated in the short or long term. Our simulations recaptured genomic patterns of several complex SVs, revealed factors affecting the formation of catastrophic SVs such as chromothripsis, and illustrated the evolutionary dynamics of ecDNAs. As it is hard to directly observe DSBs and their outcomes in practice, our in silico simulations make it possible to investigate the intermediate SV formation steps unobservable in practice, deconvolve intricacies of complex SVs, and look into the long-term evolutionary dynamics. By fitting to simulated data and single-cell whole-genome sequencing data with ABC, we demonstrated the power of our model in recovering key parameters in SV generation and revealing novel correlations among SVs. The generation of multiple SVs initiated by BFB in the same DSB repair process provides support for combining chemotherapy/radiotherapy and DNA repair inhibitors in cancer treatment [15]. The critical role of WGD and CCNE1 amplification uncovered by our model from single-cell whole-genome sequencing data also provides evidence for targeting KIF18A that is required for viability of cells with WGD or CCNE1 in the clinic [55, 56].

Most evolutionary inferences require a mutational model, such as an appropriate null model to detect driver SVs [57], and our model may serve as such a null model for hypothesis testing and model selection on related problems. By disentangling functional and non-functional SVs under selection, our model may help with distinguishing the contribution of mutation and selection in tumours driven by CIN and SVs [5]. As more SVs are anticipated to be discovered with long-read sequencing and better detection methods [10], our model will provide a useful tool to facilitate understanding of mechanisms leading to these SVs. The model may also be extended to fit other types of data such as single-cell data from experimental studies observing cell division in vitro when both CNAs and SVs can be detected in each cell [33].

To get a tractable model of complicated cell processes, we made several simplifying assumptions and hence there are a few limitations. First, we excluded biological constraints on breakpoint locations when introducing DSBs and contact probabilities when repairing DSBs. These constraints may be integrated in the future to increase the accuracy of our model, such as assigning higher DSB probabilities to fragile sites, incorporating 3D nuclear distances to derive contact probabilities, and constraining breakpoint rejoining with sequence homology as required by different repair pathways [24, 57]. Second, we simplified DNA repair and replication by directly rejoining two breakpoints and correctly duplicating all the DNAs, which prevents modelling of certain SVs resulting from HRD or complicated replication errors [25, 39]. To expand the applicability of our model to more SV types, more details such as replication-based template switching need to be incorporated. Third, the model of selection is based only on large-scale CNAs using the density of OGs and TSGs, which can be improved by considering other factors, such as SV size [34] and gene density [58], to better quantify selection strengths on SVs. Lastly, due to computational complexity, our simulations are not efficient enough to allow ABC on datasets with a large number of cells or clones, which need to be further optimized.

In conclusion, our model contributes to the development of a unified framework to investigate the biological processes underlying various patterns of CIN and to understand the relationship among related SV types. For the first time, we have been able to quantify CIN and SV generation in single-cell whole-genome sequencing data. Although these phenomena have been separately studied, we have demonstrated the close relationship of BFB cycle, chromothripsis, and ecDNA all through the processes of erroneous end joining and replication across cell cycles. Our framework provides a foundational null model for SV formation and can be used to disentangle functional and non-functional heterogeneity to guide cancer treatment development.

## 4 Methods

### Stochastic cell-cycle model of SV generation

We simulated cell growth from a single cell until reaching *N* cells with the rejection-kinetic Monte Carlo algorithm [59]. To distinguish SVs generated in a few cell cycles and SVs generated gradually over all the cell cycles, we allowed the setting of the maximum number of cycles with DSBs (*n*_*d*_). To be realistic and facilitate fitting the model to genomic data, we simulated the diploid genome with the coordinates of 22 autosomes of the reference human genome (GRCh38/hg38 by default).

The cell-cycle repair dynamics were inspired by a model of dose-dependent DSBs and rejoining after ionizing radiation [60]. In *G*_1_, we introduced *n* (either fixed or variable following Poison distribution with a pre-specified mean value *r* per cell cycle) DSBs on randomly selected chromosomes under infinite sites assumption where each DSB hits a different position. Although DSBs occur throughout the cell cycle, they mainly occur in *G*_1_ [61]. Therefore, we did not introduce additional DSBs in the other phase except in *M*. For simplicity, we excluded DSBs within telomeres and centromeres. In the analysis of chromothripsis in 634 adult tumours across 28 cancer types, the telomere and centromere regions were affected by chromothripsis in around 36% and 55% of the cases respectively [28], and hence we can still simulate a large proportion of realistic SVs. Each chromosome *c* has a probability of generating a DSB *p*_*c*_, where 1 *<*= *c <*= 22 and Σ *p*_*c*_ = 1. By default, each chromosome had the same probability of having DSBs. To facilitate the generation of chromothripsis events which often occur on one chromosome, we allowed different probabilities of DSBs on different chromosomes. This will not only generate clustered breakpoints enriched on the chromosome with the largest probability but also cause chromosome fusions derived from rejoining of inter-chromosomal breakpoints. When trying to bias towards one chromosome, we set its probability to 2/3 and the other chromosomes to have equal probability whose sum was 1/3. Once a chromosome was chosen for DSB, we randomly selected a position in a genomic interval on one homolog. As some random breakpoints may be less likely to occur in reality, we also allowed random sampling of DSBs without replacement from breakpoints extracted from detected SVs, weighted by their frequencies if available. To simulate catastrophic events such as chromothripsis, we allowed the introduction of many DSBs in the first *n*_*d*_ +1 cycle(s) and no DSBs in the subsequent cycles. When the number of divisions for a cell is larger than *n*_*d*_ (default zero, namely DSBs only occur at the first division), no more DSBs will be introduced.

We repaired a fraction (*f*_*u*_) of either all the DSBs in the cell or the new DSBs introduced in the current cell cycle with different options, where the unrepaired DSBs may be repaired in the subsequent cell cycles or not. One breakpoint will be left unrepaired when there is an odd number of breakpoints. Note that unrepaired DSBs may lead to chromosome fusion and differ from misrepaired DSBs. The first repair option is randomly joining two breakpoints, where 1*/n* of them may be faithfully rejoined on average. However, the probability of successful DSB repair often depends on the distance of the two breakpoints in the genome or the nucleus, and spatially close DSBs are more likely to be rejoined and form SVs [57, 62]. Therefore, the contact probability of two breakpoints should decrease with their genomic distance in a single chromosome or their spatial distance across chromosomes. Namely, breakpoints further away from each other on the same chromosome and breakpoints on different chromosomes are less likely to join. To be more realistic, we introduced another repair option (default), where breakpoints are rejoined based on the above observations. We first selected one breakpoint *b*_1_ and then chose the other breakpoint *b*_2_ according to the reciprocal of its distance to *b*_1_ with discrete distribution. The distance of *b*_1_ and *b*_2_ was defined as |*b*_1_ *− b*_2_| when they are on the same chromosome and 10^9^ when they are on different chromosomes. As the two breakpoints introduced by the same DSB have distance one, we assigned their contact probability to be *p*_*r*_ to control the proportion of unfaithful rejoinings [60]. The repair was based on genomic distance with a probability of correctly repairing a DSB *p*_*r*_ = 0 by default, as the correctly repaired DSBs are equivalent to no DBSs introduced. Circular DNAs with or without centromere(s), namely centric or acentric rings [60], linear chromosomes without centromere(s), and chromosome fusions may be generated due to random rejoining. For example, an isolated acentric piece without telomeres forms a circle by a new variant adjacency connecting the two ends.

In *S* and *G*_2_, we replicated the whole genome and connected broken chromosomes with lost telomeres to form fusions and di centric chromosomes, which imitates telomere crisis or shortening [63]. For circular chromosomes and complete chromosomes with both telomeres (e.g. pink chromosome in Fig. 1), we made an exact copy. Due to unrepaired breakpoints, some chromosomes may lose one or both telomeres. For a non-circular chromosome lacking one telomere or both telomeres (e.g. purple and green chromosomes in Fig. 1), we linked it to its copy, imitating sister chromatid fusion. The genomic segment lacking both telomeres will form a circular DNA. We consider the genomic segment lacking both telomeres and a centromere as an ecDNA, which can be formed either during *G*_1_ or during chromosome fusion.

In *M*, we distributed the replicated chromosomes into two daughter cells according to the number of centromeres *N*_*c*_ in each chromosome. If *N*_*c*_ = 1, the chromosome was inherited by either daughter cell. If *N*_*c*_ = 0, the chromosome was randomly distributed to a daughter cell. As a result, one daughter cell may have two copies of this chromosome and the other daughter cell may lose it completely, leading to reciprocal gain and loss. If *N*_*c*_ *>* 1, the chromosome was subject to DSB, imitating chromosome segregation errors such as bridge breakage. Previous experiments suggest that direct bridge breakage generated either simple break or local fragmentation which may affect two or more chromosomes [20]. We simulated a simple break by breaking the chromosome at feasible breakpoints according to the locations of all the centromeres in the fused chromosome to ensure that each derivative chromosome had a single centromere and was allocated to a different daughter cell when *N*_*c*_ = 2 and to a random daughter cell when *N*_*c*_ *>* 2. We simulated local fragmentation by breaking the derivative chromosome with one centromere after a simple break into *n*_*l*_ pieces, where *n*_*l*_ followed a Poison distribution with a pre-specified mean *m*_*l*_, and each piece was randomly allocated to a daughter cell. When *n*_*l*_ *>* 0, we introduced local fragmentation with a probability of 0.5 to avoid too much shattering. We also simulated WGD in each cell with probability *p*_*w*_, assuming at most one WGD in a cell, by allocating all the chromosomes to one daughter cell and letting the other daughter cell die.

We used OG-TSG score to measure the fitness of a cell according to the density of OGs and TSGs on chromosome-level or arm-level (default) CNAs [2]. We computed the survival probability of each cell with the formula [64]:

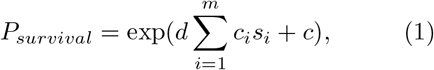

where *d* = 0.00039047, *c* = *−*0.036132164, *m* is the number of chromosomes or arms, *c*_*i*_ is the average total copy number of chromosome or arm *i*, and *s*_*i*_ is the corresponding OG-TSG score. When the genome is normal, *P*_*survival*_ = 0.9689715. Assuming exponential growth of cells, the population size at time *t* is

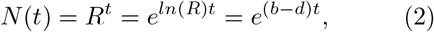

where *R* is the average number of offspring of a cell. Suppose 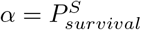 is the probability for a daughter cell to survive, where *S >* 0 is introduced to control the magnitude of selective pressure [58], then the offspring distribution is *P* = (*p*_0_, *p*_1_, *p*_2_), where *p*_0_ = (1 *− α*)^2^, *p*_1_ = *α*(1 *− α*), and *p*_2_ = *α*^2^. Therefore,

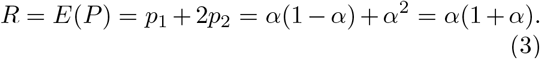

Then we can get the relationship between the survival probability of a cell *α*and its birth/death rate:

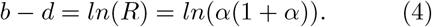

Assuming the birth and death rate of the normal cell and its daughter cell are *b*_0_, *d*_0_, *b*_1_, and *d*_1_ respectively, then

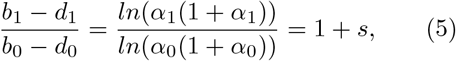

where *s* is the selection coefficient measuring the strength of natural selection on the daughter cell.

When *s* = 0, it means neutral evolution and the daughter cell has the same birth/death rate as the parent cell. When *s >* 0, it means the daughter cell is positively selected and has either a higher birth rate or a lower death rate. When *s <* 0, it means the daughter cell is negatively selected and has either a lower birth rate or a higher death rate. We then derived the value of *s* from *α*_0_ and *α*_1_ and updated the birth/death rate of the daughter cell accordingly. Note that when *S* is no larger than 15, the larger the *S*, the stronger the selection is.

The program generated detailed output files for inference and validation, which include SVs and haplotype-specific CNAs in each cell, the derivative genome represented by a list of connected intervals for each cell, and the cell lineage relationship generated in the branching process. The CNAs were reported at both segment level of variable size and bin level of fixed size with pre-specified locations. We obtained haplotype-specific copy numbers for a segment by merging consecutive intervals with the same copy number on a homolog of a chromosome. To get haplotype-specific copy numbers for non-interval adjacencies, we traversed all the adjacencies and grouped them by positions and orientations. To get bin-level copy numbers which are often detected in bins of fixed size such as 500 Kbp, we allocated each segment to bins of size *B*. We computed bin IDs *b*_*i*_ for a segment *i* with start position *s*_*i*_ and end position *e*_*i*_ by computing the start bin ID 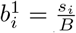 and the end bin ID 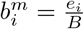, where 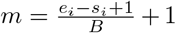 is the number of bins. For the two bins at the ends of a segment *i*, there may be other segments covering these bins, so the contribution of segment *i* to the copy number of these bins is weighted by the overlap size.

We computed multiple summary statistics for each run of the simulation (Table S5), among which the fraction of cells with WGD, the fraction of different types of SVs (deletion, duplication, inversions, and intra-chromosomal SVs), the frequency distribution of breakpoints present in different numbers of cells, the percentage of genome altered (PGA), and the mean and standard deviation of pairwise divergence were used for inference with ABC. PGA and the mean and standard deviation of pairwise divergence were computed based on total and haplotype-specific copy numbers, respectively. A bin is altered if its copy number is different from the ploidy, which is two for normal cells and four for cells with WGD when considering total copy number, and one for normal cells and two for cells with WGD when considering haplotype-specific copy number. We defined PGA as the percentage of bins that are altered (in any cell) from the ploidy across all cells and pairwise divergence between two cells as the proportion of bins with different copy numbers that are altered from the ploidy.

### Graph representation of the genome

We defined a breakpoint *b* as the position on a chromosome *c* (1 *<*= *c <*= 22) of size *l* (0 *< b < l*), with a label *H ∈ {A, B}*, where *A* and *B* represent the two homologs of *c* that are inherited from each parent. A DSB at position *j* of a chromosome introduced two breakpoints *j* and *j* + 1, each of which was represented as a node in the graph. Each breakpoint divided a segment [*i, k*] into two intervals [*i, j*] and [*j* + 1, *k*]. Following conventions [27], the orientation of the left break-point *j* was denoted by +/HEAD (the right side or head of interval [*i, j*]), whereas the orientation of the right breakpoint *j* + 1 was denoted by -/TAIL (the left side or tail of interval [*j* + 1, *k*]). Most breakpoints were introduced in *G*_1_ as a result of interphase DSBs, and the remaining breakpoints were generated in *M* due to bridge breakage. The adjacency orientations on the same chromosome, determined based on the involved breakpoints, indicate simple SV-like patterns of four types: tandem duplication-like (*−/*+), deletion-like (+*/−*), head-to-head inversion (+*/*+), and tail-to-tail inversion (*−/−*), whereas adjacencies across two different chromosomes indicate inter-chromosomal SVs.

The genome was represented as a set of paths in the graph *G*, where a path is a walk in *G* alternating between interval and reference/variant adjacency edges. A path represents a derivative chromosome and may contain a varying number of centromeres and telomeres (no telomere, left/right telomere, and both telomeres). We represented a circular path by adding one more adjacency between the two ends of the corresponding linear path. We did a walk along the graph at the end of *G*_1_ to get all the paths in the genome before DNA replication. Among the adjacencies introduced in one cell cycle, most occurred during breakpoint rejoining in *G*_1_ and the remaining variant adjacencies happened during chromosome fusion in *S* to represent head-to-head and tail-to-tail inversions. In *M*, due to bridge breakage, some genomic intervals may get broken and lead to new adjacencies in *G*_1_ of the next cell cycle.

### Generation and analysis of simulated data

To reduce visualization burden and get a variety of SVs in Fig. 1c, we only introduced five DSBs at each cell cycle, which were biased towards chr1.

To learn the effect of a single BFB with simple breaks, we introduced one DSB on a random chromosome at the first cell cycle and left it unrepaired until 100 cells under both neutral evolution and selection (selection strength *S* = 5). To further show our model can simulate more complex patterns such as chromothripsis under local fragmentation, we set the mean number of DSBs during local fragmentation to 50 to get more breakpoints in a short time. We introduced two DSBs at the first cell cycle with 50% of DSBs correctly repaired (*p*_*r*_ = 0.5) at each cell cycle, which left one unrepaired DSB after the first cell cycle. This unrepaired DSB led to ongoing BFB cycles, where the resultant bridges underwent simple breaks or local fragmentation randomly in each cell cycle. We used the same seed for simulations under neutral evolution and selection so that the resultant differences were due to selection rather than stochasticity.

To simulate chromoplexy, we introduced 10 misrepaired DSBs per cycle on random chromosomes in each cell cycle under neutral evolution. To see the parallel evolution of haplotype-specific CNAs, we continued the simulation until *N* = 200.

For simulations on the formation of chromothripsis, we simulated data with DSB rate per cycle *r* ∈ {10, 30}, fraction of unrepaired DSBs per cycle *f*_*u*_ ∈ {0, 0.1, 0.3}, and mean number of DSBs per fragmentation) *n*_*l*_ ∈ {0, 10, 30} (50 datasets for each parameter setting) under neutral evolution. As chromothripsis often occurs on one chromosome in a short time, we simulated for one or two cycles and biased DSBs towards chr1 (as in Fig. 1c). We only introduced interphase DSBs at the first cycle for some simulations with two cycles to compare with the effects of two consecutive cycles with interphase DSBs.

For parameter inference on simulated data, we set *S* = 1 with GRCh37/hg19 as a reference, since cancer genomes often underwent selection and genome sequencing data were often analysed by mapping to GRCh37/hg19. We did not model local fragmentation to reduce the number of parameters to estimate. To get realistic break-points, we sampled from breakpoints of a primary prostate cancer sample (ID: 2340225) in COSMIC [65], which has the most (14,626) SVs. We simulated until 10 cells, as it is computationally expensive to do ABC on a larger population size. We simulated 180 datasets with DSB rate per cycle *r ∈ {*10, 30*}*, fraction of unrepaired DSBs per cycle *f*_*u*_ *∈ {*0.1, 0.3, 0.5*}*, and probability of WGD per cell *p*_*w*_ *∈ {*0.1, 0.3, 0.5*}*, with 10 datasets for each parameter setting.

We computed the overlap of simulated CNAs with cancer-related genes from COSMIC Cancer Mutation Census [65]. We used ShatterSeek [27] to detect chromothripsis from simulated SVs and CNAs. We counted the numbers of high-confidence chromothripsis events which have at least six interleaved intra-chromosomal SVs and at least seven segments with oscillating copy number states. Some of the simulated amplifications may be located in ecDNAs, which will cause higher-level amplifications and violation of the criteria of chromothripsis and lead to fewer detected chromothripsis events. We detected seismic amplification with the pipeline in [8] and chromoplexy with Jabba [7]. Although most amplifications were located in ecDNAs, they can be integrated into the genome through circular recombination, which was not included here for simplicity. Hence, we treated ecDNAs as homogeneously staining regions that are reintegrated into the chromosome when detecting seismic amplification.

### Analysis of single-cell whole-genome sequencing data

Three mutational signatures were detected in the PDX datasets in [43], where 8 datasets belong to HRD-Dup (homologous recombination deficiency-duplication) tumours which are enriched in BRCA1 mutations and small tandem duplications, 3 datasets belong to tumours which are enriched in large tandem duplications and CDK12 mutations, and 12 datasets belong to FBI tumours that were used for our inference. As we assumed correct DNA replication other than chromosome fusion and no sequence homology when repairing DSBs to reduce complexity, our model is not suitable for SVs which were probably generated from HRD or other replication errors [25, 39]. Therefore, we excluded 11 PDX datasets belonging to HRD-Dup and TD tumours and used the 20 remaining datasets for the subsequent analysis. We computed the empirical DSB rates for each dataset as the average number of subclonal SVs per clone expansion, namely the number of sub-clonal SVs divided by the number of clones minus 1.

For inference with ABC, we simulated data with GRCh37/hg19 as a reference under selection (*S* = 1) without local fragmentation. The fractions of breakpoints in telomere or centromere regions for all the datasets are less than 20% (Table S6), so the exclusions of DSBs on these regions did not affect the inference. Because some datasets share clonal SVs and CNAs, which should be presented in the common ancestor of these clones, we started the simulation with the clonal SVs. To reduce stochasticity, the positions of new DSBs were sampled from the detected breakpoints based on their clonal prevalence, where break-points present in more clones were more likely to be sampled.

Note that the observed breakpoints or SVs are the results of both mutation and selection, which may suffer from false positives and false negatives due to the challenges in accurate detection [4]. In addition, not all the detected SVs arise from erroneous DNA repair and replication, such as SVs caused by transposable element insertion or virus integration. Therefore, the parameters inferred from real data only provided approximate estimates to better understand the underlying mutational processes.

### Running ABC SMC on simulated data and single-cell whole-genome sequencing data

We used the function ABCSMC in the Julia package ApproxBayes [59] to run the ABC SMC algorithm. In the algorithm, a set of sampled parameter values is propagated through a series of intermediate distributions (populations) with decreasing tolerances until the target tolerance *ϵ* is reached, where each sampled parameter value gets a weight based on its importance. We took 500 samples from each population to estimate the posterior distributions of parameters and set the maximum number of simulations to 1,000,000. We mainly varied the prior distributions of parameters and target tolerance and set the other parameters to the default values. For consistency, we assumed that the parameter priors were uniform for all the inferences on both simulated data and single-cell whole-genome sequencing data. The prior distributions of fraction of unrepaired DSBs and probability of WGD followed uniform distribution [0, 1]. For the inference on simulated data, we assumed the prior distribution of DSB rate followed uniform distribution [0, *M*], where the upper limit *M* was 50, and set *ϵ* to 0.2, which was sufficient to get accurate results in a reasonable amount of time. For the inference on single-cell whole-genome sequencing data, we assumed the prior distribution of DSB rate followed uniform distribution [0, *M*], where the upper limit *M* was 50 for datasets with empirical DSB rate less than 50 and 100 otherwise, and set *ϵ* to 0.5 because it would take a much longer time for convergence with a smaller *ϵ*. For 13 datasets with wider posterior distributions in Fig. 6a, ABC SMC stopped at a higher *ϵ* because the last *ϵ* was within 5% of the previous population.

## 5 Data availability

The simulated data and processed single-cell whole-genome sequencing data are available at Zenodo (https://doi.org/10.5281/zenodo.10114638).

## 6 Code availability

Our model is available at https://github.com/_ucl-cssb/CIN_SV. The package ApproxBayes used for ABC SMC is available at https://github.com/marcjwilliams1/ApproxBayes.jl.

### Supplementary information

Supplementary tables and figures.

## Acknowledgments

The authors acknowledge the use of the UCL Myriad High Throughput Computing Facility (Myriad@UCL) and the UCL Department of Computer Science High Performance Computing Cluster, and associated support services, in the completion of this work.

### Appendix A Extended Data

**Fig. A1.**
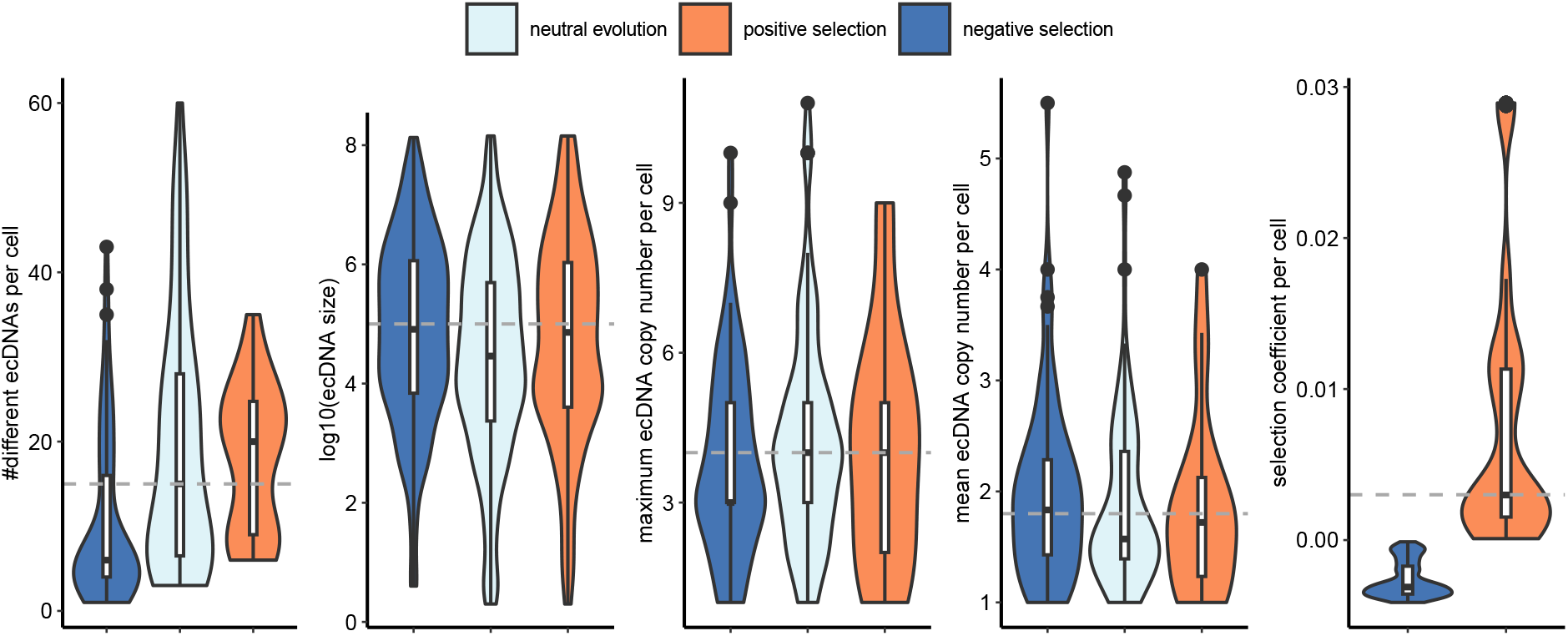
The distributions of summary statistics of ecDNAs from the simulation with local fragmentation (Fig. 2c). There are 73, 95, and 22 cells under negative selection, neutral evolution, and positive selection, respectively. The dashed lines are used as references for comparison. The box plots show the median (centre), 1st (lower hinge), and 3rd (upper hinge) quartiles of the data; the whiskers extend to 1.5 times of the interquartile range (distance between the 1st and 3rd quartiles); data beyond the interquartile range are plotted individually.

**Fig. A2.**
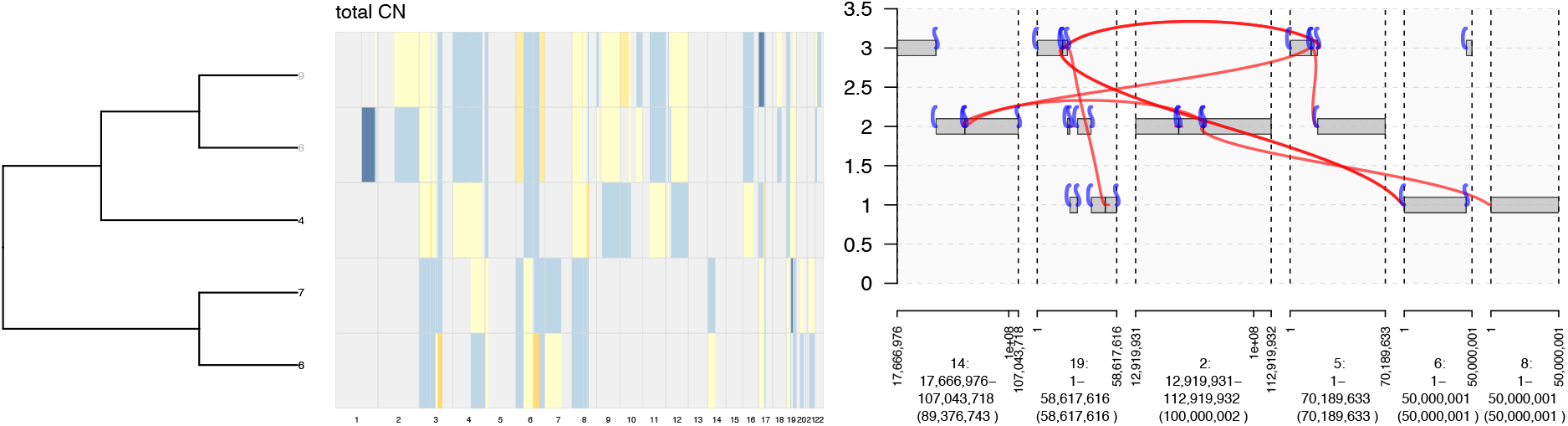
The cell lineage tree of five cells and their corresponding copy number heatmap across all chromosomes from the simulation with multiple misrepaired DSBs, as well as a detected chromoplexy in cell 6. The cells without chromoplexy are shown in grey.

**Fig. A3.**
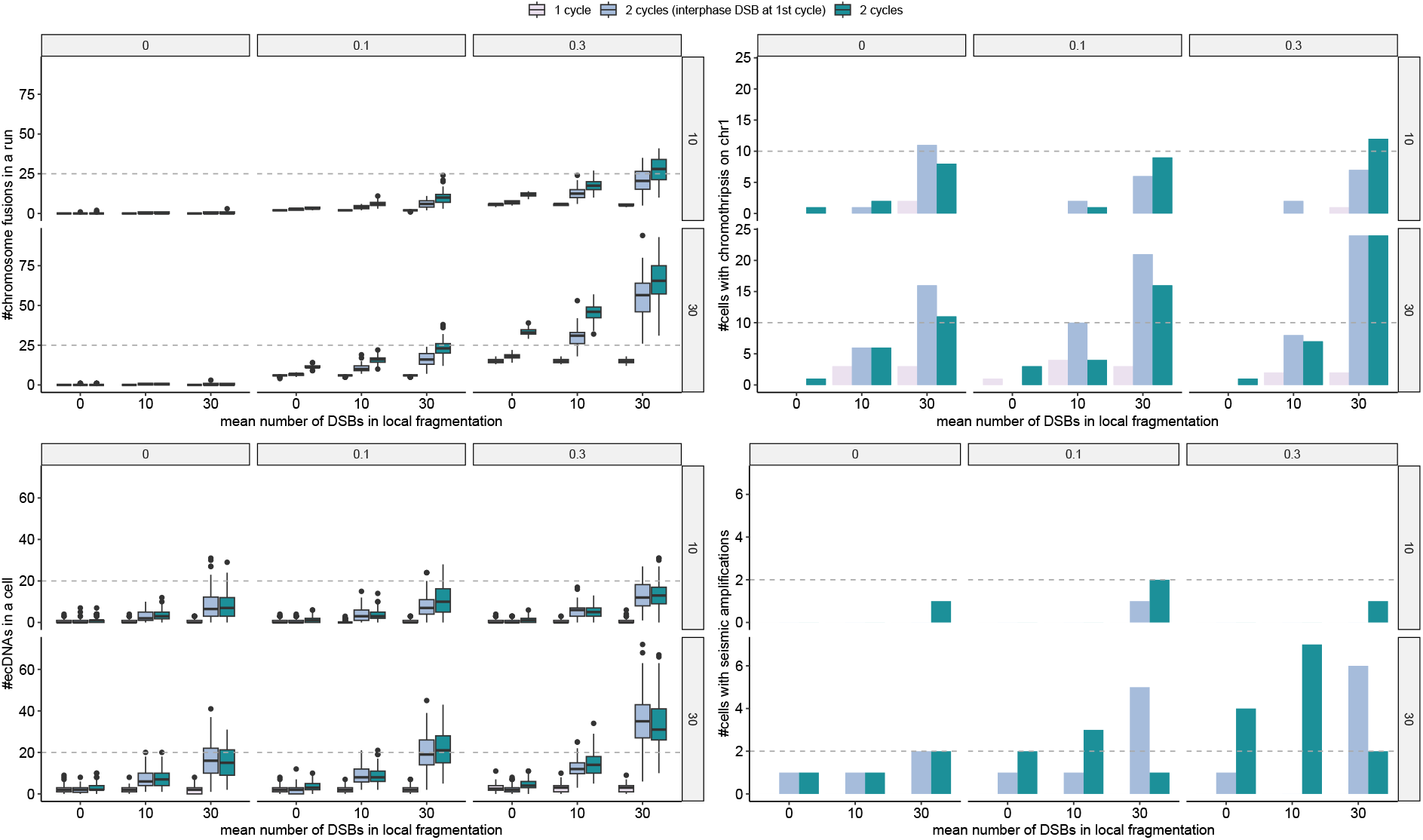
The distributions of complex SVs under different parameter settings in one or two cell cycles shown in groups different from those in Fig. 3. The labels at the top of each plot indicate fraction of unrepaired DSBs per cycle and the labels at the right indicate DSB rates per cycle. For each run, we consider the two cells at the end of either the first or second cycle depending on the number of cycles. The box plots show the median (centre), 1st (lower hinge), and 3rd (upper hinge) quartiles of the data; the whiskers extend to 1.5 times of the interquartile range (distance between the 1st and 3rd quartiles); data beyond the interquartile range are plotted individually.

**Fig. A4.**
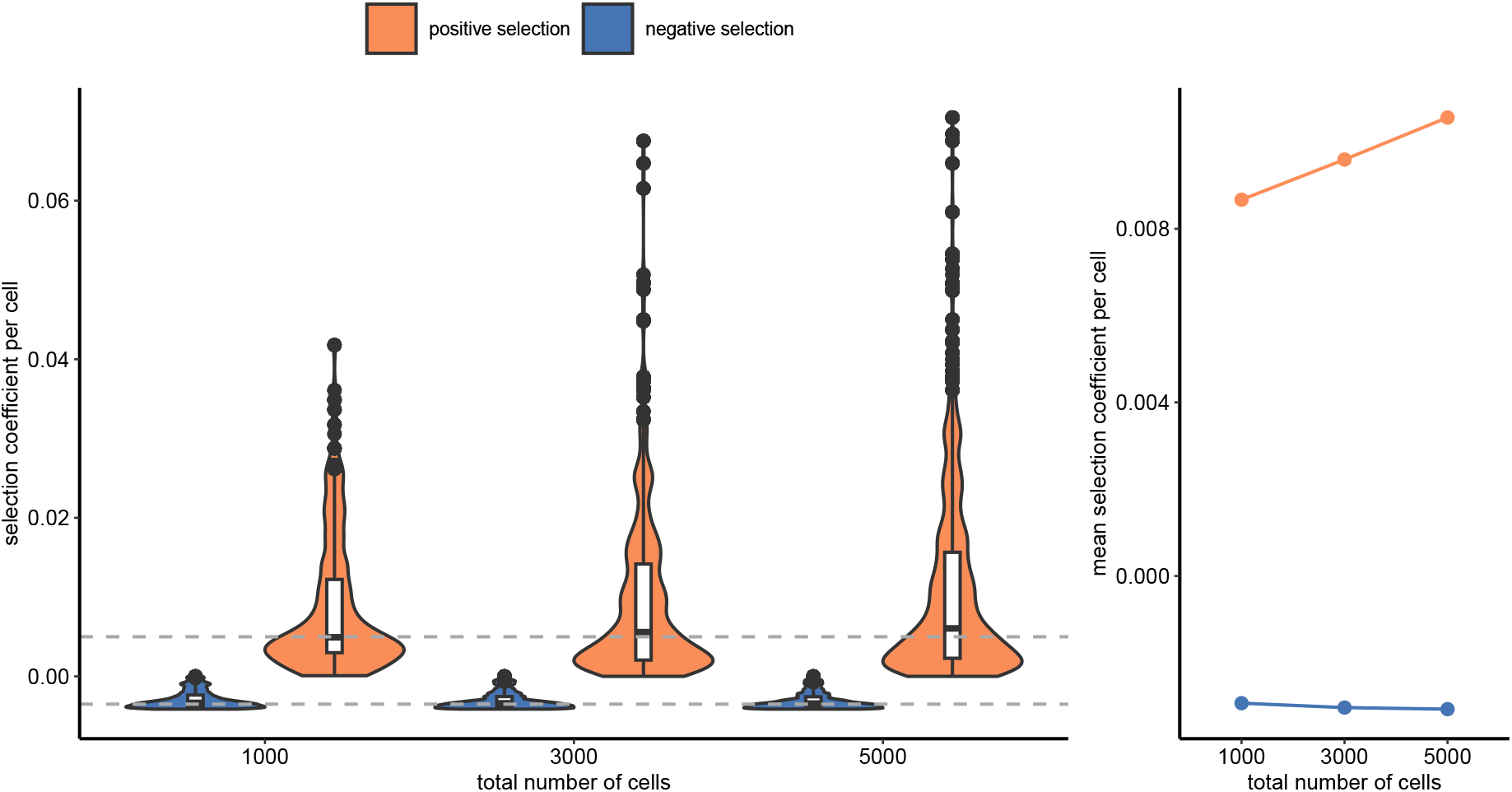
The distributions of selection coefficients of cells (Fig. 4) under selection over time. The box plots show the median (centre), 1st (lower hinge), and 3rd (upper hinge) quartiles of the data; the whiskers extend to 1.5 times of the interquartile range (distance between the 1st and 3rd quartiles); data beyond the interquartile range are plotted individually.

**Fig. A5.**
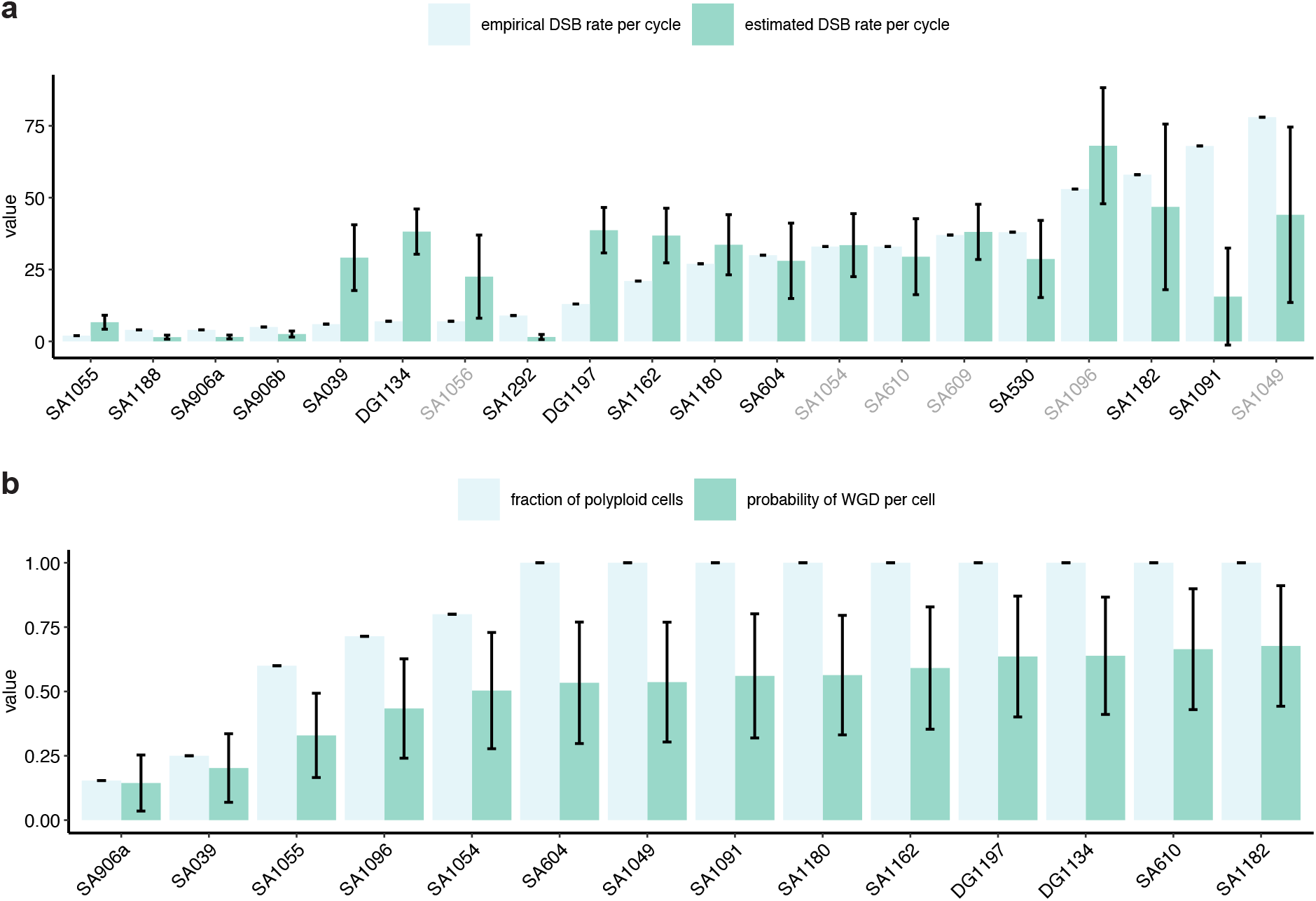
The comparison of estimated parameters with empirical values in single-cell whole-genome sequencing datasets. The error bar for the estimated parameter represents the weighted standard deviation of the posterior means. The names of datasets with WGD are shown in grey.

**Fig. A6.**
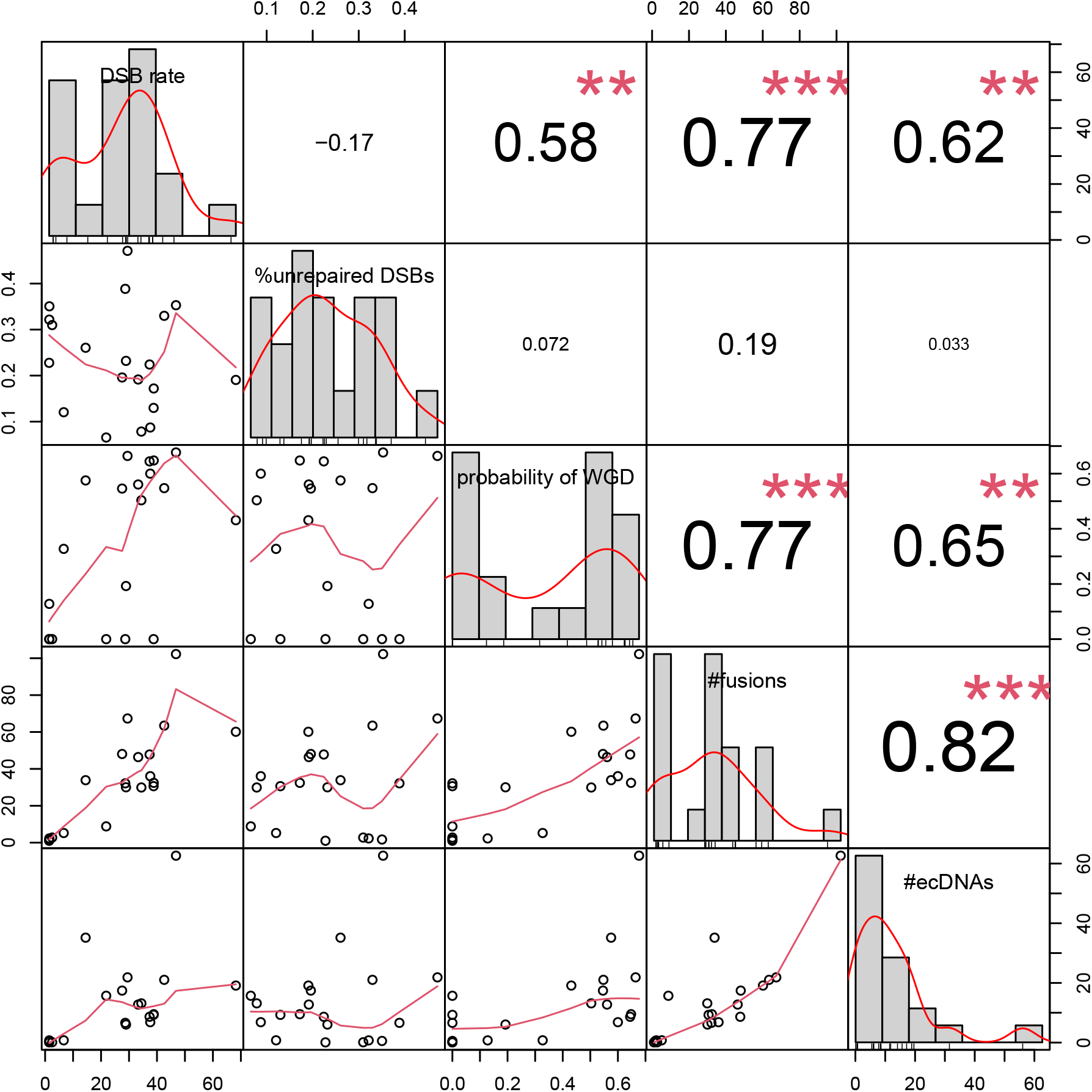
The pairwise correlations of parameters inferred from single-cell whole-genome sequencing datasets. The parameters include the inferred DSB rate per cycle, fraction of unrepaired DSBs per cycle, probability of WGD per cell, mean number of fusions per cycle, and mean number of ecDNAs per cell. The Spearman correlation coefficient and corresponding significance are shown for each pair of parameters. The indicator of significance: * – p-val *<* 0.05, ** – p-val *<* 0.01, *** – p-val *<* 0.001.

**Fig. A7.**
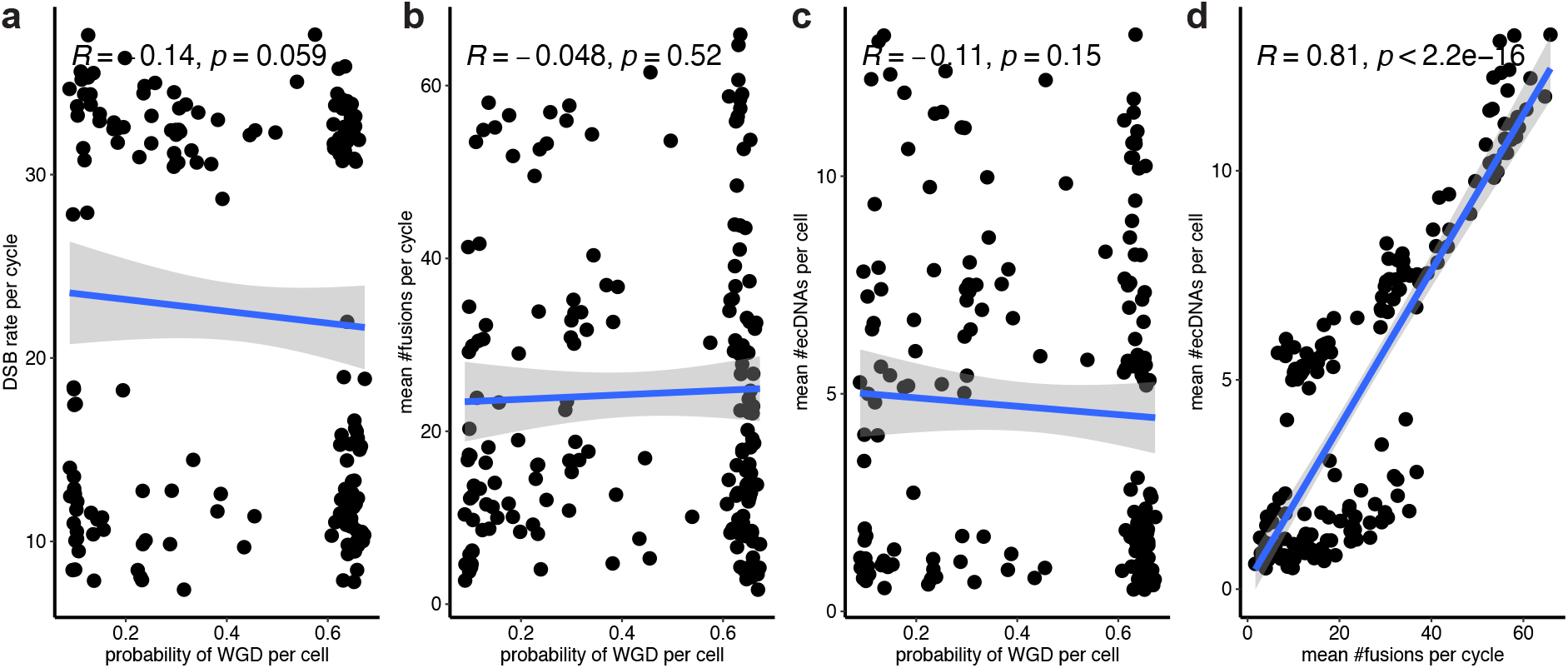
The pairwise correlations of parameters inferred from simulated data (Fig. 5). The Spearman correlation coefficient and corresponding p-value are shown for each pairwise plot. The shaded area shows the 95% confidence interval of linear regression. There are 180 data points on each plot, where the same set of parameters generated 10 data points.

**Fig. A8.**
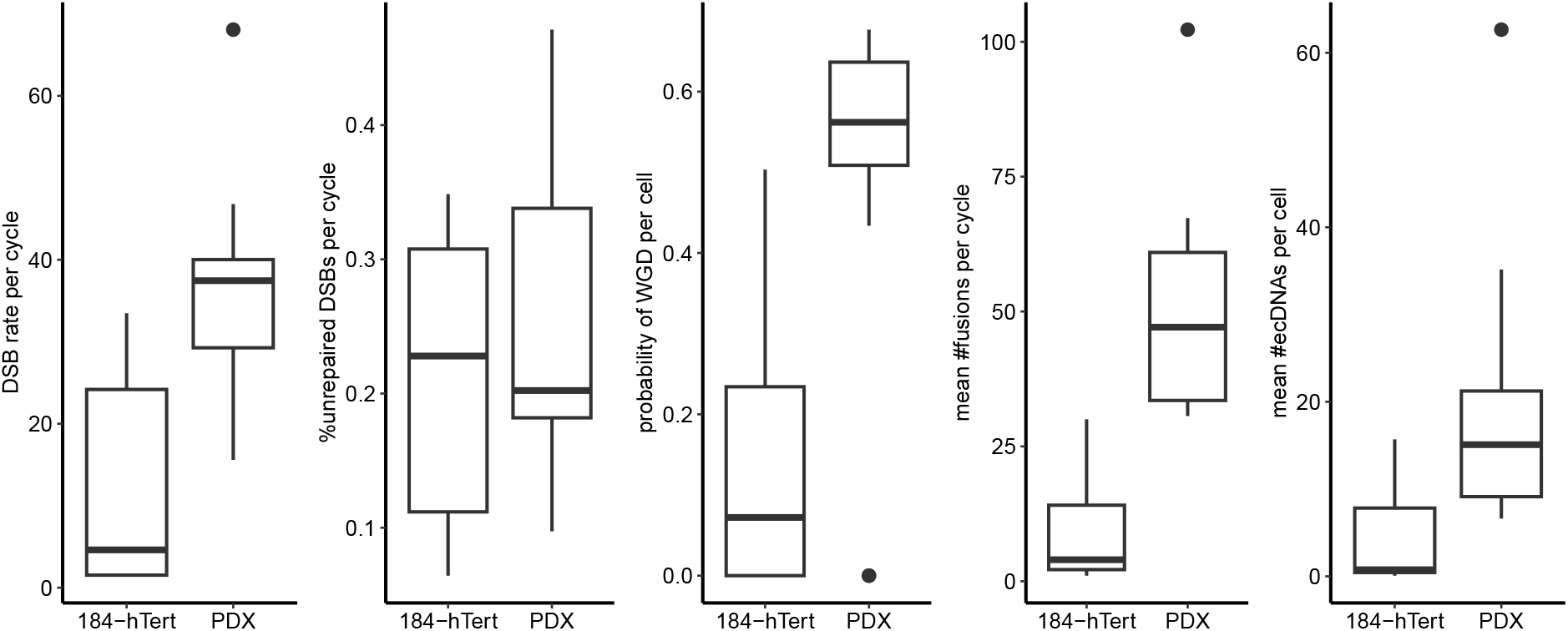
The comparison of posterior means of parameters estimated from single-cell whole-genome sequencing datasets of different cell types. The box plots show the median (centre), 1st (lower hinge), and 3rd (upper hinge) quartiles of the data; the whiskers extend to 1.5 times of the interquartile range (distance between the 1st and 3rd quartiles); data beyond the interquartile range are plotted individually.

