## Supplementary File for "Cell-cycle dependent DNA repair and replication unifies patterns of chromosome instability"

### Supplementary document for Cell-cycle dependent DNA repair and replication unifies patterns of chromosome instability

Table S1: The major input parameters of the stochastic cell-cycle model.

| **parameter** | **symbol** | **default value** |
|---|---|---|
| DSB rate per cycle | $r$ | 5 |
| fraction of unrepaired DSBs per cycle | $f_u$ | 0 |
| probability of WGD per cell | $p_w$ | 0 |
| population size (number of cells) | $N$ | 2 |
| birth rate per cycle | $b$ | 1 |
| death rate per cycle | $d$ | 0 |
| number of DSBs per cycle | $n$ | 0 |
| maximum ID of cycles with DSBs | $n_d$ | 0 |
| mode of repairing | $m_p$ | 1 (based on distance) |
| probability of correct DSB repair | $p_r$ | 0 |
| mean number of DSBs during local fragmentation | $n_l$ | 0 |
| probability of DSB on a chromosome $c$ | $p_c$ | 1 / 22 |
| model of evolution | $m$ | 0 (neutral) |
| strength of selection | $S$ | 1 |

Table S2: The total copy numbers of oncogenes (OGs) and tumour suppressor genes (TSGs) overlapping with SVs in simulated data with simple breaks (Fig. 2a).

| **type** | **name** | **total copy number** | **#cells (neutral model)** | **#cells (selection model)** |
|---|---|---|---|---|
| **OG** | CARD11 | 3 | 3 | 5 |
| | | 5 | 4 | 6 |
| | | 7 | 3 | 0 |
| | | 9 | 0 | 3 |
| | EGFR | 3 | 7 | 10 |
| | | 4 | 3 | 3 |
| | | 5 | 0 | 2 |
| | RAC1 | 3 | 3 | 5 |
| | | 5 | 4 | 6 |
| | | 7 | 3 | 0 |
| | | 9 | 0 | 3 |
| **TSG** | PMS2 | 1 | 86 | 81 |
| | SFRP4 | 1 | 86 | 87 |

Table S3: The numbers of complex SVs detected in simulated data with local fragmentation (Fig. 2c).

| **model** | **#chromothripsis (cycle appearing)** | **#ecDNAs (cycle appearing)** | **#seismic amplifications (cycle appearing)** | **#cells with chromothripsis, seismic amplification, and ecDNAs** |
|---|---|---|---|---|
| **neutral** | 37 (4) | 95 (4) | 12 (5) | 3 |
| **selection** | 36 (5) | 95 (3) | 8 (5) | 1 |

Table S4: The total copy numbers of oncogenes (OGs) and tumour suppressor genes (TSGs) involved in genome rearrangements in simulated data with local fragmentation (Fig. 2c).

| **type** | **name** | **total copy number** | **#cells (neutral model)** | **#cells (selection model)** |
|---|---|---|---|---|
| OG | ACVR1 | 3 | 10 | 9 |
| | | 5 | 10 | 7 |
| | | 7 | 3 | 3 |
| | | 9 | 0 | 3 |
| | | 11 | 3 | 0 |
| | CTNNA2 | 3 | 2 | 2 |
| | | 5 | 6 | 3 |
| | | 7 | 2 | 0 |
| | | 11 | 0 | 2 |
| | | 17 | 2 | 0 |
| | | 23 | 0 | 2 |
| | CXCR4 | 3 | 11 | 12 |
| | | 5 | 6 | 4 |
| | | 7 | 0 | 2 |
| | | 13 | 2 | 0 |
| | IDH1 | 3 | 9 | 4 |
| | | 5 | 2 | 2 |
| | | 7 | 3 | 0 |
| | MYCN | 3 | 2 | 2 |
| | | 5 | 6 | 3 |
| | | 7 | 2 | 0 |
| | | 11 | 0 | 2 |
| | | 17 | 2 | 0 |
| | | 23 | 0 | 2 |
| | SF3B1 | 3 | 3 | 6 |
| | | 5 | 9 | 3 |
| | | 7 | 3 | 2 |
| | SIX2 | 3 | 2 | 2 |
| | | 5 | 6 | 3 |
| | | 7 | 2 | 0 |
| | | 11 | 0 | 2 |
| | | 17 | 2 | 0 |
| | | 23 | 0 | 2 |
| | XPO1 | 3 | 2 | 2 |
| | | 5 | 6 | 3 |
| | | 7 | 2 | 0 |
| | | 11 | 0 | 2 |
| | | 17 | 2 | 0 |
| | | 23 | 0 | 2 |
| TSG | ACVR2A | 1 | 68 | 68 |
| | ASXL2 | 1 | 85 | 89 |
| | BARD1 | 1 | 78 | 87 |
| | CASP8 | 1 | 80 | 87 |
| | DNMT3A | 1 | 85 | 89 |
| | LRP1B | 1 | 73 | 73 |
| | MSH2 | 1 | 85 | 89 |
| | MSH6 | 1 | 85 | 89 |
| | TMEM127 | 1 | 64 | 76 |

Table S5: The major summary statistics generated by the stochastic cell-cycle model.

| **summary statistics** | **range** | **level** | **status in inference** |
|---|---|---|---|
| number of ecDNAs | $[0, \infty]$ | per cell | not used |
| number of chromosome fusions | $[0, \infty]$ | per cycle | not used |
| percentage of genome altered (PGA) | [0,1] | across all cells | used |
| mean and standard deviation of pairwise divergence | [0,1] | across all cells | used |
| frequency distribution of breakpoints | [0,1] | across all cells | used |
| fraction of cells with WGD | [0,1] | across all cells | used |
| fraction of different types of SVs | [0,1] | across all cells | used |

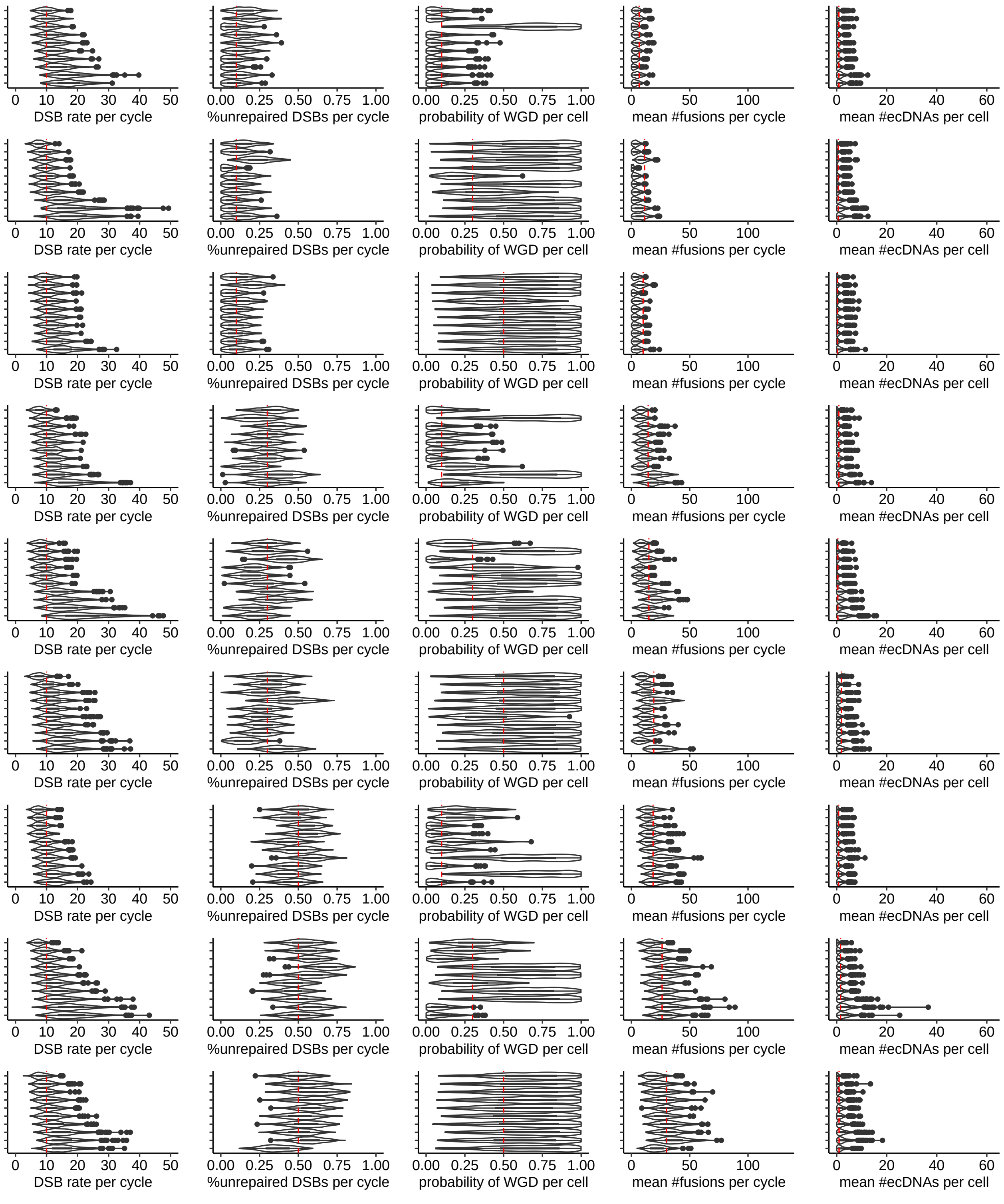


Fig. S1: **The posterior distributions of three inferred parameters and the posterior predictive distributions of two summary statistics from simulated data with DSB rate per cycle $r = 10$.** Each row represents the results for one set of parameters $(r, f_u, p_w)$, where $f_u$ is fraction of unrepaired DSBs per cycle and $p_w$ is probability of WGD per cell. The red dasheds indicate the true parameter values. The results are sorted by the inferred values of $r$. The violin plots for the three inferred parameters are weighted.

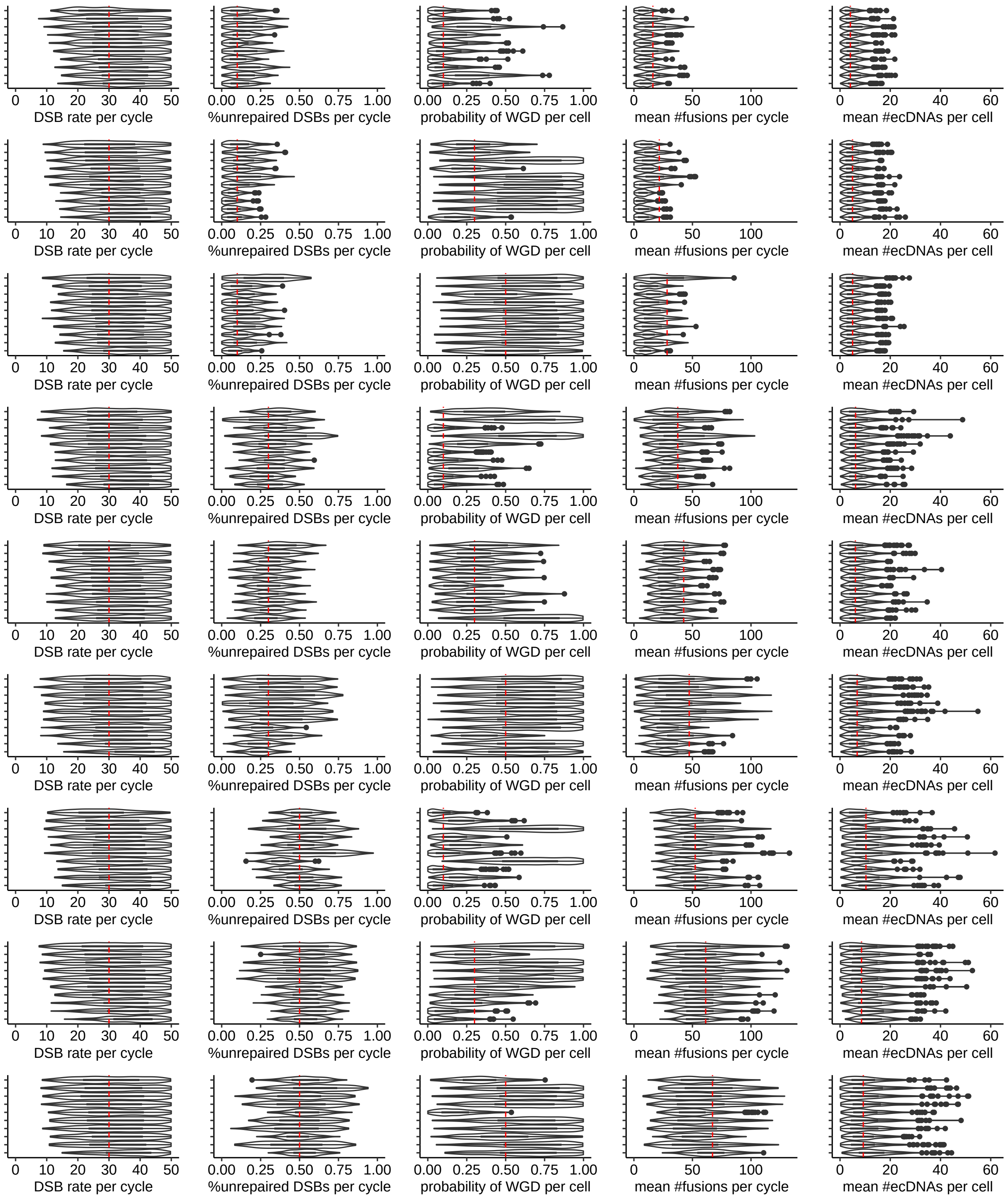


Fig. S2: **The posterior distributions of three inferred parameters and the posterior predictive distributions of two summary statistics from simulated data with DSB rate per cycle $r = 30$.** Each row represents the results for one set of parameters $(r, f_u, p_w)$, where $f_u$ is fraction of unrepaired DSBs per cycle and $p_w$ is probability of WGD per cell. The red dashed lines indicate the true parameter values. The results are sorted by the inferred values of $r$. The violin plots for the three inferred parameters are weighted.

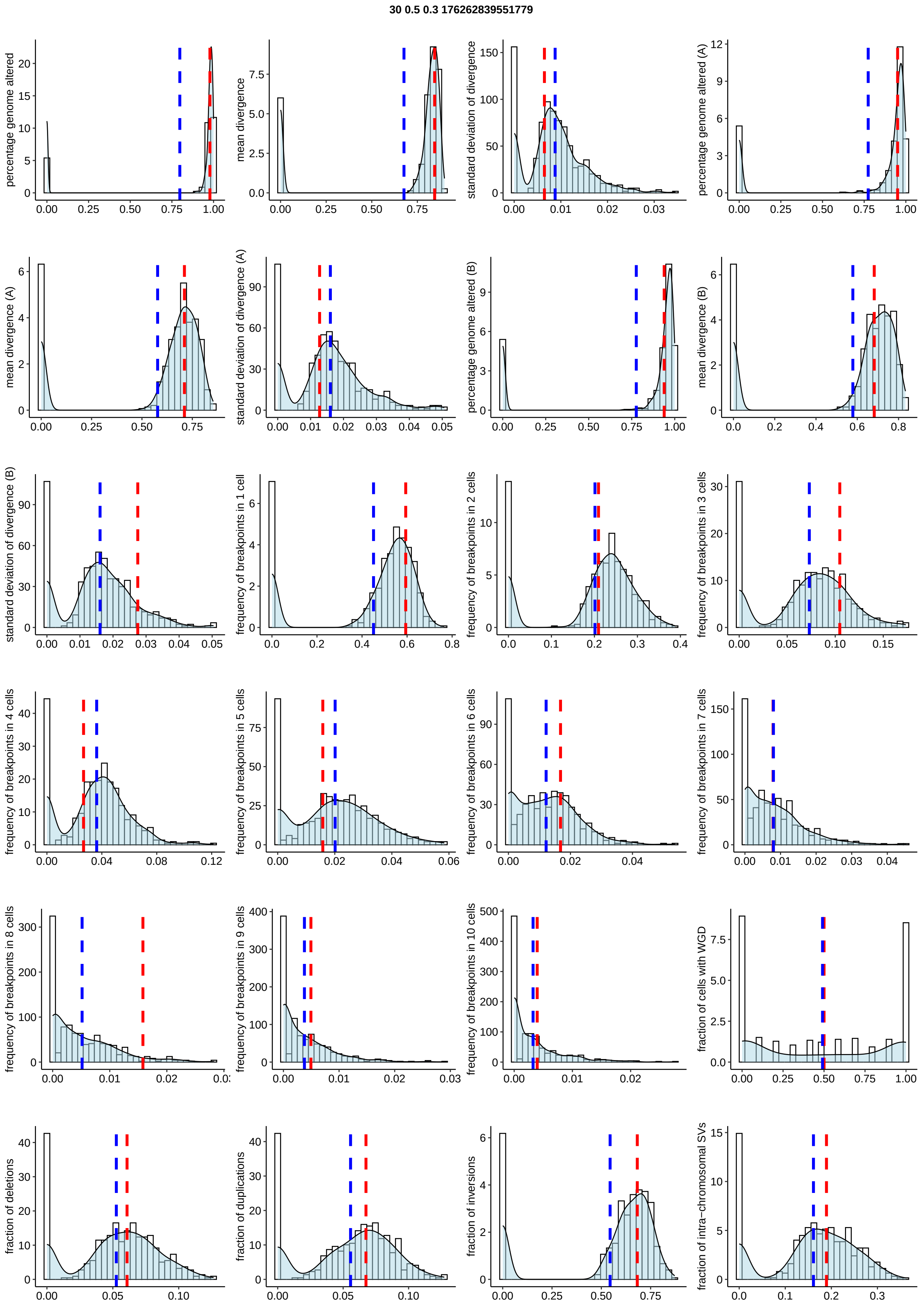


Fig. S3: **The posterior predictive distributions of summary statistics used in inference from a simulated dataset.** The red dashed lines indicate the observed values. The blue dashed lines indicate the posterior means. The title indicates the true parameter values used for simulating the data, where DSB rate per cycle $r = 30$, fraction of unrepaired DSBs per cycle $f_u = 0.5$, probability of WGD per cell $p_w = 0.3$, and the random seed is 176262839551779.

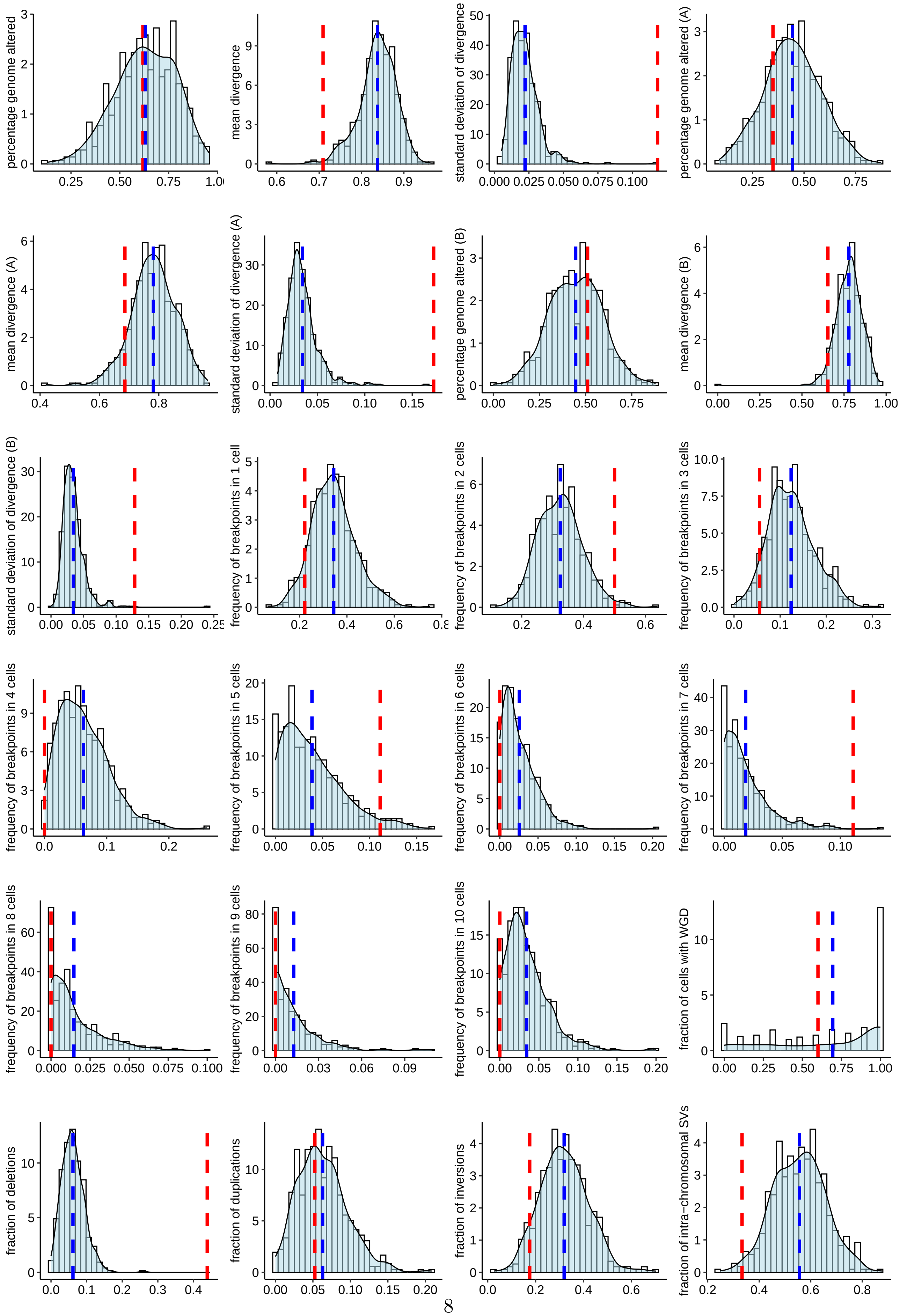




Fig. S4: **The posterior predictive distribution of summary statistics used in inference from a single-cell whole-genome sequencing dataset SA1055.** The red dashed lines indicate the observed values. The blue dashed lines indicate the posterior means.
